# Exploring Gene Expression Patterns and Evolutionary Responses in Host-Parasite inteactions: Insights from the *Schistocephalus solidus* – Threespine stickleback System

**DOI:** 10.1101/2023.07.18.547692

**Authors:** Anika M. Wohlleben, Javier F. Tabima, Néva P. Meyer, Natalie C. Steinel

**Affiliations:** Insitute of Zoology and Evolutionary Research, Friedrich Schiller University Jena, Jena, Germany; Biology Department, Clark University, Worcester, Massachusetts, USA; Department of Biological Sciences, University of Massachusetts – Lowell, Lowell, Massachusetts, USA

## Abstract

Helminth parasites pose a significant threat to host survival and reproductive success, imposing strong selection pressure on hosts to evolve countermessures (e.g., immune responses and behavioral changes). To gain insights into the underlying mechanisms of host-parasite co-evolution, we examined differences in gene expression in immune tissues of two Alaskan stickleback (*Gasterosteus aculeatus*) populations with varying susceptibility to infection by the cestode *Schistocephalus solidus*. Our analyses revealed distinct patterns of immune gene expression at the population-level in response to infection. Infected fish from the high infection population displayed signs of immune manipulation by the parasite, whereas this phenomenon was absent in fish from the low infection population. Notably, we found significant differences in immune gene expression between the populations, with uninfected Rocky Lake fish showing up-regulation of innate immune genes associated with inflammation compared to uninfected Walby Lake fish. These findings highlight the divergent evolutionary paths taken by different stickleback populations in their response to the same parasite.

## Introduction

As helminths are frequent parasites in natural fish populations (Holmes, 1990; Kennedy, 2009; Lizama et al., 2004), there should be strong selection pressure for hosts to develop countermeasures to suppress/avoid infection (Summers et al., 2003). Host-parasite dynamics can drive host immunocompetence and infection resistance on the one hand (Eizaguirre et al., 2011), and parasite virulence and infection success on the other (Decaestecker et al., 2007). To study host-parasite co-evolution and population-level differences in parasite susceptibility we used the threespine stickleback–*Schistocephalus solidus* system. When oceanic stickleback invaded freshwater habitats after the last glacial maximum, they encountered the cestode *S. solidus* for the first time (Simmonds and Barber, 2016), as it is absent in marine environments. The independent colonization of freshwater bodies has generated multiple, independent, natural, co-evolutionary experiments that we can use to answer fundamental questions about host adaptations to novel parasites and how host-parasite interactions are shaped by an ever-changing environment. Freshwater stickleback populations exhibit differences in their susceptibility and prevalence of *S. solidus* infection (Heins and Baker, 2003; Weber et al., 2017a). The mechanisms underlying these population-level differences are highly variable, but it is evident that the host immune response plays a crucial role. The difference in host-parasite co-evolution between populations highlights the importance of studying the complex dynamics of these interactions on a multi-population scale.

Both the stickleback innate and adaptive immunity are manipulated by *S. solidus* upon infection (Scharsack et al., 2004a, 2007; Wohlleben et al., 2018). Differences in gene expression related to ROS production and MHC have been observed between populations with high and low infection prevalence (Lohman et al., 2017b). Fibrosis, a recently described anti-parasite response in stickleback that is shown to reduce *S. solidus* size (Hund et al., 2022; Weber et al., 2022), is also highly host population-specific (Hund et al., 2022; Weber et al., 2022). Assessment of stickleback immune variaton has predominantly been performed in lab-reared fish. However, to gain a comprehensive understanding of immune functions and phenotypes of free-living populations, it is essential to study wild organisms (Martin et al., 2011; Pedersen and Babayan, 2011). Therefore, to identify mechanisms underlying differences in *S. solidus* susceptibility among host populations, we characterized the immune profiles of infected and uninfected stickleback from two Alaskan populations that differ in *S. solidus* infection prevalence: Walby Lake (high infection, low fibrosis) and Rocky Lake (low infection, high fibrosis).

To assess habitat-specific and local variation in immune response, we focused on transcriptomes in natural-infected, wild-caught stickleback. Environmental factors strongly influence immune expression patterns (Abolins et al., 2017; Robertson et al., 2016; Stutz et al., 2015), and studying naturally infected fish allowed us to account for these environmental factors (e.g. genetic diversity of host and parasite) which are often underrepresented in laboratory-based studies. Here we show that immune gene expression profiles differ between naturally infected fish from two stickleback populations that differ in their susceptibility to *S. solidus* infections.

## Methods

### Study lake selection and fish trapping

To detect differences in immune gene expression between stickleback populations with variable *S. solidu*s prevalence we analyzed gene expression in two stickleback populations from the Matanuska-Susitna valley of southcentral Alaska. Walby Lake (61° 37’ 9"N, 149° 12’ 39"W) has high *S. solidus* infection prevalence, with infection rates between 20% and 80% (Heins et al., 2010). Rocky Lake (61° 33’ 23"N, 149° 49’ 29"W) in contrast, has low infection prevalence, typically ranging between 2% and 11% (Wohlleben, 2022). Both lakes have similar ecological properties with the lakes being surrounded by small houses and birch and spruce trees. On both lakes, birds like loons, the terminal host for *S. solidus*, are common visitors (John Baker, personal communication).

In the first week of June 2021 (Rocky Lake: June 11^th^; Walby lake June 12^th^), we caught stickleback via unbaited, 6 mm wire-mesh minnow traps set overnight near shore. On the day of capture, we euthanized the fish with MS-222 (Syndel, Ferndale, WA, USA) and immediately dissected the head kidneys, a primary immune organ in teleost fish (Geven and Klaren, 2017; Press and Evensen, 1999), which we stored in RNAlater (Thermo Fisher Scientific, Waltham, MA, USA). We screened each fish for macroscopic parasites, and recorded fibrosis presence/absence scores, standard length (body length from the tip of the nose to the last vertebrae), weight, and stickleback sex. Rocky Lake fish displayed a high level of fibrosis and fibrotic fish had encapsulated the parasite within fibrotic tissue. We stored head kidneys at -80 °C until RNA extraction. Collections and procedures were approved by annual Aquatic Resource Permits from the Alaskan Department of Fish and Game (SF2021-106), and IACUC approved animal care protocols (033R).

### RNA extraction

We extracted RNA from the collected head kidneys using the RNeasy Mini Kit (Qiagen, Hilden, Germany). After thawing on ice, we removed the head kidneys from RNAlater and placed them in a tube with ∼10 ceramic beads (1.4 mm) and 600 µL of RTL Buffer enriched with 6 µL β-mercaptoethanol (Sigma-Aldrich, St. Louis, MO, USA) to avoid sample degradation by RNases. We next used a Mini-Beadbeater 8 (0 – 2800 SPM: BioSpec Products, Bartlesville, OK, USA) to lyse the sample. Subsequently, we followed the manufacturers protocol for RNA extraction (Qiagen RNeasy Mini Kit). Each RNA sample was precipitated in 5 µL of 3 M sodium acetate and 125 µL of 100% ethanol. After incubation at -20°C for 16– 18 hours, we centrifuged samples with a Sorvall Legend Micro 21R Microcentrifuge (Thermo Fisher Scientific, Waltham, MA, USA) at 21,000 x g at 4°C for 15 minutes. We discarded the supernatant and added 125 µL 70% ethanol before centrifugation at 21,000 x g at 4°C for 3 minutes. We discarded the supernatant, and air dried the RNA pellets for 40–60 minutes before resuspending the RNA in nuclease-free water. We removed DNA from the samples with the Thermo Scientific RapidOut DNA Removal Kit (Thermo Scientific, Waltham, MA, USA) following the manufacturers protocol before storing the RNA samples at -80°C. We assessed RNA quality via Nanodrop and RNA concentration via Qubit (RNA quantification, broad Range) (Thermo Scientific, Waltham, MA, USA), to adjust sample concentration to ∼ 30 ng/µl.

### Tag-Seq and Bioinformatics

For Walby Lake stickleback, we analyzed transcriptomes of 10 infected and 10 uninfected fish. For Rocky Lake stickleback, we analyzed transcriptomes of 9 uninfected fish, 9 infected fish with encysted *S. solidus,* and 8 infected fish with non-encysted *S. solidus*. Library prep and TaqSeq were performed at the Genomic Sequencing and Analysis Facility (GSAF) at the University of Texas, Austin. TagSeq libraries were sequenced on the Illumina NovaSeq 6000 SR100 system with standard coverage (between 3,000,000 and 5,000,000 reads per sample) as single reads. We roughly followed a previously described iRNAseq pipeline, (https://github.com/z0on/tag-based_RNAseq) (Lohman et al., 2016; Meyer et al., 2011). First, we assessed QC scores and removed duplicate reads and trimmed adapters using the cutadapt package (version 4.1) (Martin, 2011). Subsequently we aligned the sequenced transcripts to the reference genome, using Bowtie 2 (version 2.4.5) (Langmead and Salzberg, 2012). We then aligned reads to the ensembl *Gasterosteus aculeatus* genome (BROAD S1, May 2022) (http://ftp.ensembl.org/pub/release-107/fasta/gasterosteus_aculeatus/cdna/). The percentage of mapped reads was between 50% and 60% (Table S1).

### Differential gene expression and GO term analysis

To assess differential gene expression, we generated read counts per gene using the samcount command (Bowtie 2 version 2.4.5) (Langmead and Salzberg, 2012) and used the R statistical software package DESeq2 (Love et al., 2014). For Rocky Lake stickleback we included fish sex as factor in the DESeq2 analysis as we found several gene co-expression networks to be correlated with it. We did not find a correlation between gene networks and fish sex in Walby Lake fish and hence did not include it in the DESeq2 analysis. We filtered transcripts to remove rows with only few reads (13) and defined a log2-fold change of 0 as equal expression (Lohman et al., 2017a). We multiply test corrected all p-values using a 10% false discovery rate (FDR) (Benjamini and Hochberg, 1995). After ordering differentially expressed genes based on the smallest p-values, we extracted a FASTA file. We extracted Gene Ontology (GO) terms from the FASTA file via the InterproScan package (Jones et al., 2014).

### Weighted correlation network analysis (WGCNA)

We normalized raw read counts via variance stabilizing transformation (VST) (Anders and Huber, 2010) before plugging them into the R statistical software package WGCNA (Langfelder and Horvath, 2008). For our signed networks, we used a soft thresholding power of 24, and a minimum module size of 15 genes, and merged modules with greater than 80% similarity. We used the software Cytoscape (version 3.9.1) (Shannon et al., 2003) to visualize the gene networks within each module. We retrieved gene annotations from the Zebrafish Information Network (ZFIN), University of Oregon, Eugene, OR 97403-5274; http://zfin.org/; [September 2022] and collected data on protein sequences associated with genes that were not found in the Zebrafish Information Network from The UniProt Consortium (The UniProt Consortium, 2019). If no protein sequences were found, we used the ensemble database to look for orthologs (Cunningham et al., 2022).

We created all graphs for this study in R studio (v. 4.2.1; R Core Team, 2022) or Cytoscape (Shannon et al., 2003) and edited them in Inkscape (Inkscape Project, 2020) and GIMP (v. 2.10.22; The GIMP development Team, 2019). All R packages are listed in Table 1.

**Table 1:** R packages used for statistical analysis and figure creation.

| R-package | Used for... |
| --- | --- |
| xlsx (Dragulescu and Arendt, 2020) | Importing excel files into R |
| DESeq2 (Love et al., 2014) | Assessing differential gene expression |
| tidyverse (Wickham et al., 2019) | Chaining commands |
| ggplot2 (Wickham 2016) | Plot creation – Data visualization |
| Seqinr (Charif and Lobry, 2007) | Generating & extracting FASTA file with annotations |
| WGCNA (Langfelder and Horvath, 2008) | Weighted correlation network analysis |
| RCy3 (Gustavsen et al. 2019) | Visualizing and analyzing gene network in Cytoscape using R |
| GO.db (Carlson 2019) | Separating GO terms into categories for visualization |

## Results

### Immune gene expression in Walby Lake stickleback, a high infection population

To identify genes that may contribute to parasite susceptibility in a high infection stickleback population, we compared TagSeq results between uninfected and *S. solidus*-infected Walby Lake stickleback. TagSeq reads produced 12,789 genes for Walby Lake samples, which is comparable to other studies of this nature (Fuess et al., 2021b; Lohman et al., 2017a). Out of those 12,789 genes, 7.27% were down-regulated and 6.58% were up-regulated in infected compared to uninfected stickleback (Wald, p < 0.1 after 10% FDR correction). Out of the top 100 differentially expressed genes in Walby Lake stickleback, 73 were up-regulated and only 27 were significantly down-regulated. GO enrichment analysis showed that differentially expressed genes were enriched for proteolysis, integral membrane proteins, protein binding and ATP binding. We found few genes (29) to be associated with the GO term of immune response (Figure 1). However, upon closer examination of individual differentially expressed genes, we discovered that many of them were associated with innate and adaptive immune responses, despite not being identified through the GO term analysis, and instead, several of these genes were associated with other GO terms. For instance, *lck* (*lymphocyte-specific protein tyrosine kinase*) and *zap-70* (*zeta-chain-associated protein kinase 70*), which play crucial roles in T cell receptor signaling in mammals, were significantly down-regulated in infected Walby stickleback (Table 2). Although these genes are linked to other GO terms such as “ATP binding”, they were not listed under the “Immune Response” term. Similarly, *cd40* (Table 2), a protein essential for T and B cell activation was listed under “Protein binding” but not “Immune Response”.

**Figure 1:**
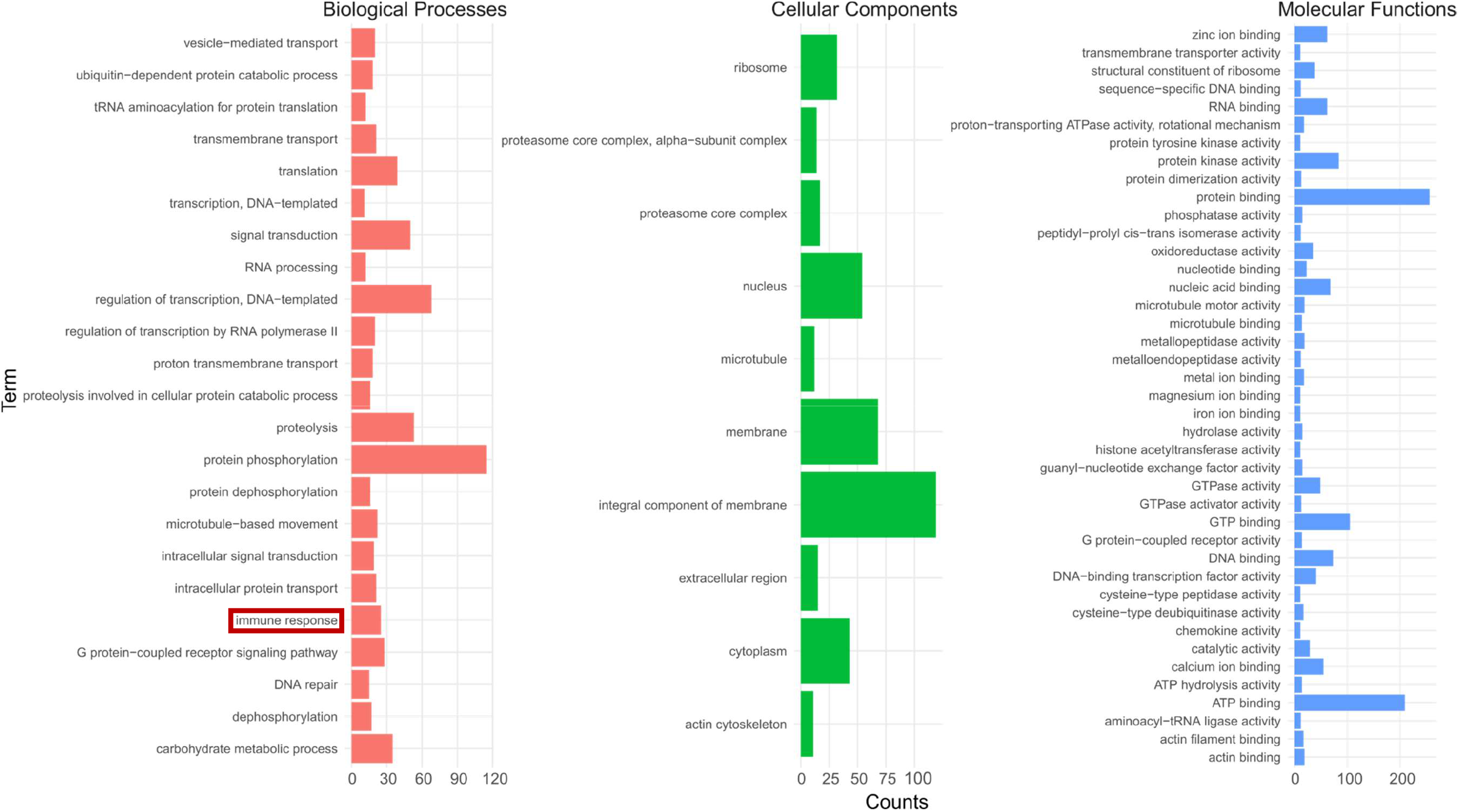
Gene Ontology (GO) term enrichment analysis for Walby Lake stickleback for both up- and down-regulated genes. Go terms of the categories Biological processes, Cellular Components, and Molecular Functions are shown in red, green, and blue, respectively. The red box highlights the Go term of Immune Response.

**Table 2:**
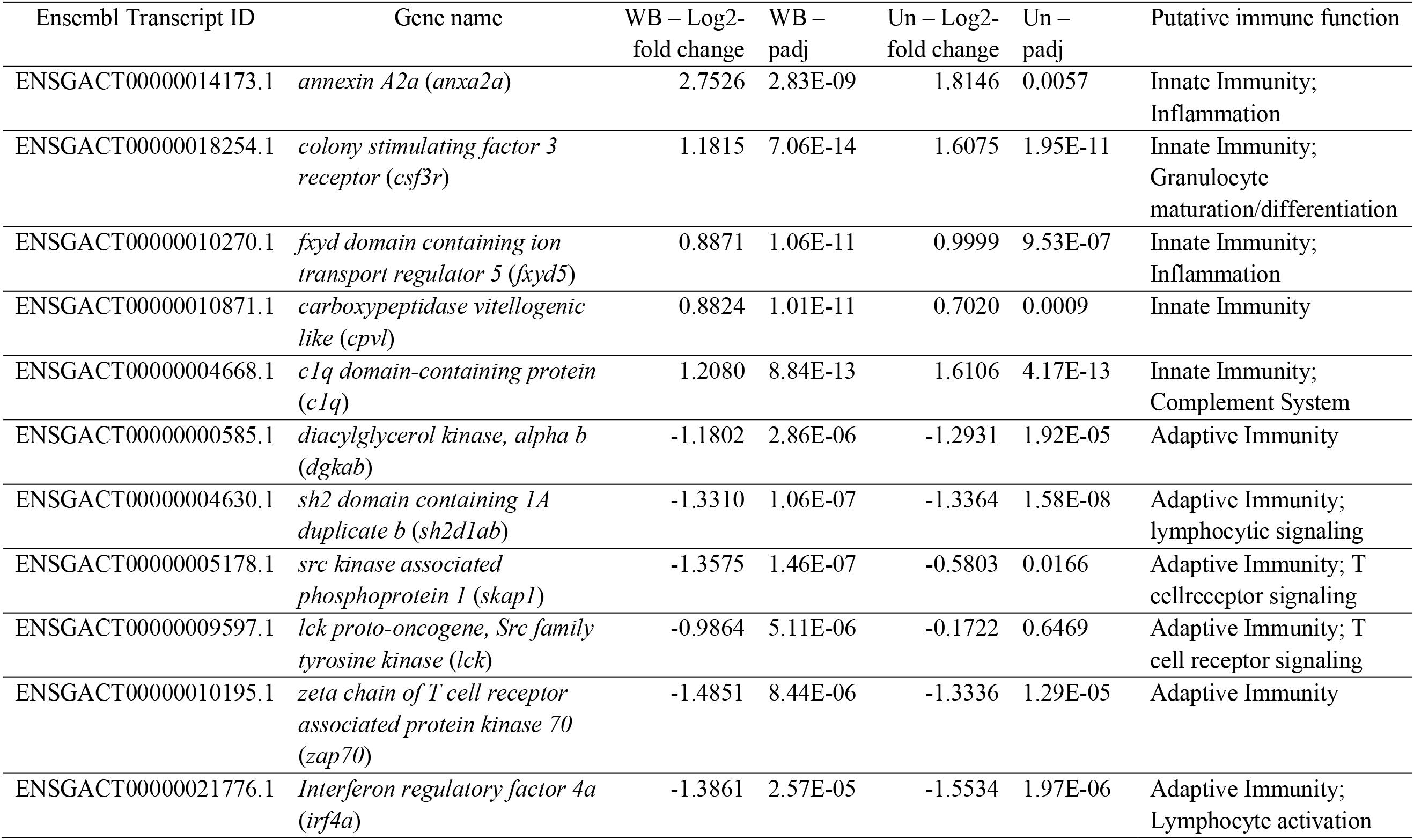
Selection of top differentially expressed genes associated with immune functions in infected vs. uninfected Walby Lake (WB) stickleback and the same genes as a function of population (Rocky Lake vs. Walby Lake) in uninfected stickleback (Un). The log 2-fold change refers to the differences in infected compared to uninfected fish. The p-value (padj) is adjusted for multiple testing via Benjamini-Hochberg (10% FDR).

Interestingly, many innate immune genes were up-regulated in infected fish (Table 2, Table 3). For example, *cpvl* (*carboxypeptidase vitellogenic like*), a serine carboxypeptidase, which is proposed to be involved in the inflammatory protease cascade and the digestion of phagocytosed particles in the lysosome, was up-regulated. We also found that *anxa2a (annexin a2a*) and its receptor *s100a10* were both up-regulated in infected Walby Lake stickleback (Table 2). *annexin* is involved in several immune pathways, including the modulation of reactive oxygen species (He et al., 2016, p. 20), the regulation of the inflammatory response (triggered by *tlr4*) (Zhang et al., 2015) and the complement cascade (Martin et al., 2012). One of the functions of the complement system is to bridge the innate and adaptive immune system via the complement component 1q (c1q). c1q can directly bind to the surface of certain pathogens or it can bind to antibody:antigen complexes, thus linking the innate and adaptive immunity (Bennett et al., 2017; Nakao et al., 2011).

**Table 3:**
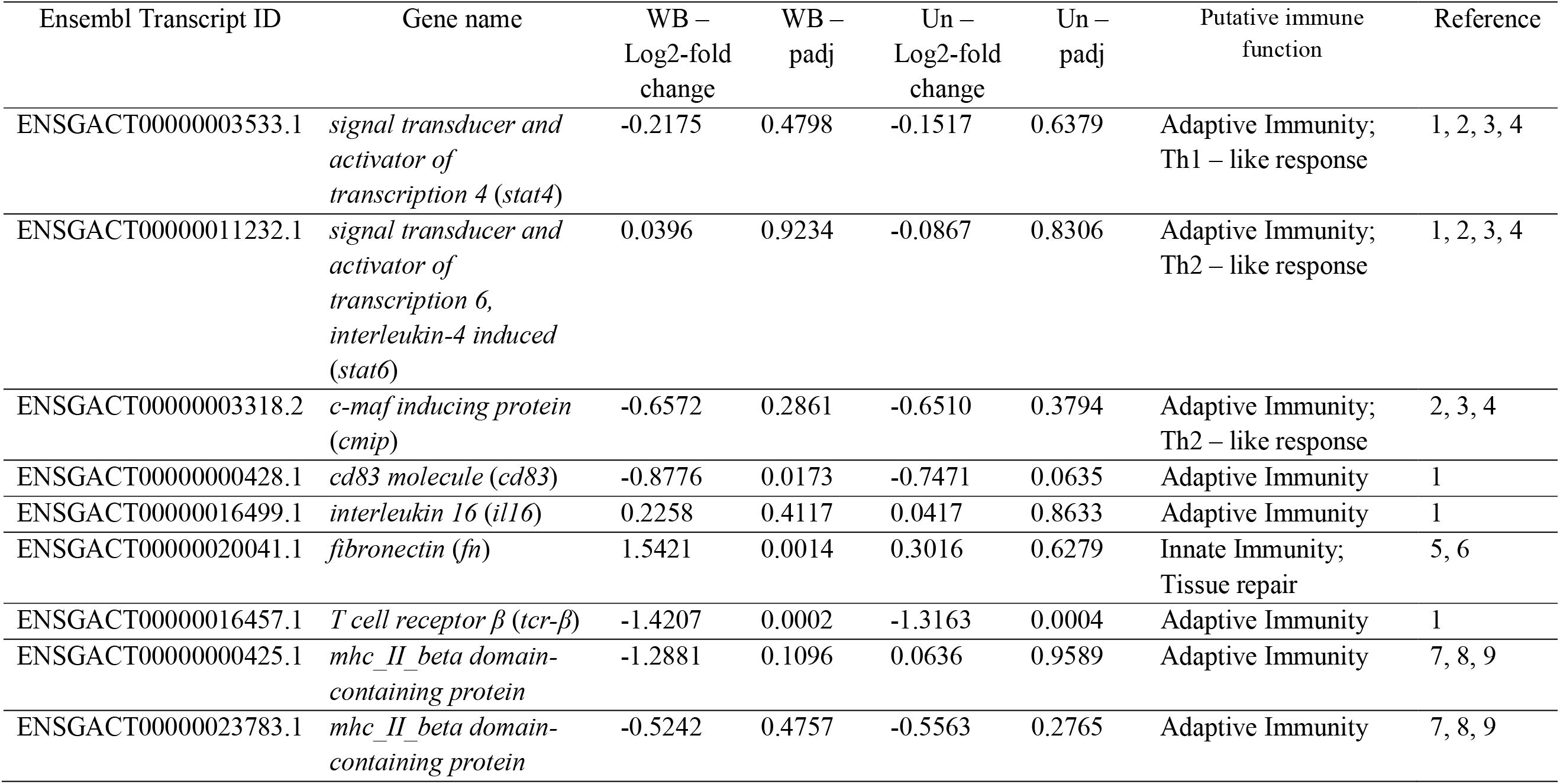
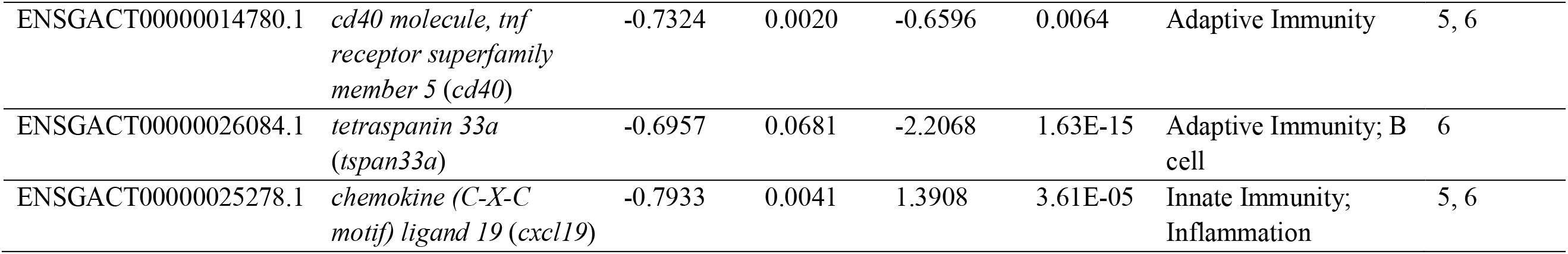
List of candidate genes in infected vs. uninfected Walby Lake (WB) stickleback and in uninfected stickleback from both populations (Un). The log 2-fold change refers to the differences in infected compared to uninfected fish in WB and the difference between uninfected Rocky compared to Walby Lake fish in Un. The p-value (padj) is adjusted for multiple testing via Benjamini-Hochberg (10% FDR). Key for References: 1)(Piecyk et al., 2019) 2) (Robertson et al., 2016) 3) (Whiting et al., 2018) 4) (Whiting et al., 2020) 5) (Fuess et al., 2021c) 6) (Lohman et al., 2017) 7)(Haase et al., 2016) 8) (Kurtz et al., 2004) 9) (Wegner et al., 2006)

In contrast to the up-regulation of innate immune genes, we found many genes involved in the adaptive immune response to be down-regulated with infection. For example, the beforehand mentioned *lck* and *zap-70* (Table 2), which are important for T cell receptor signaling; T cells (i.e. cytotoxic, helper, and regulatory) are important players in the adaptive immune response. Their roles include directly killing infected host cells, activating other immune cells, and producing cytokines. We also observed a significant down-regulation of *sh2d1ab* (*sh2 domain containing 1A duplicate b*), which is expressed in T cells, natural killer cells, and some B cells, where it modulates signal transduction pathways (reviewed in (Veillette et al., 2008). Interestingly, the expression of *cmip* (*c-maf inducing protein)*, which codes for an adaptor protein that inhibits the activation of *lck* (Oniszczuk et al., 2020), was not affected by infection status.

When comparing our results with those from previous transcriptomic studies, we observed moderate overlap between expression patterns (Table 3). For example, we found *cd40*, a co-stimulatory protein that is expressed by antigen-presenting cells and required for T and B cell activation (Grewal and Flavell, 1996; Munroe and Bishop, 2007) to be down-regulated, which is in line with previous reports (Lohman et al., 2017a). Further, we found the previously discussed *anxa2a* to be up-regulated, similar to previous work by Fuess and colleagues (Fuess et al., 2021c) and Lohman and colleagues (Lohman et al., 2017a). In contrast, while we did not find differences in expression of the transcriptional activators *stat4* (Th1 response) and *stat6* (Th2 response), previous work found both to be differentially expressed in response to *S. solidus* infection (Piecyk et al., 2021, 2019). Similarly, a marker for B cell activation, *tspan33a* (*tetraspanin 33a*), was not differentially expressed in the present study but found to be up-regulated in response to infection in the past (Lohman et al., 2017a).

### Weighted correlation network analysis for Walby Lake stickleback

To determine correlation patterns among gene sets (modules), as well as between gene sets and sample features (e.g. infection status, host sex) for Walby Lake stickleback, we analyzed the RNAseq data via WGCNA (weighted correlation network analysis). We identified nine co-expression modules (Figure 2), several of which were correlated with infection status and number of *S. solidus* per fish. The infection status was positively correlated with the black module and negatively correlated with the brown and grey modules. The number of parasites per host was positively correlated with the red module and negatively correlated with the grey and brown modules. None of the modules was associated with stickleback sex.

**Figure 2:**
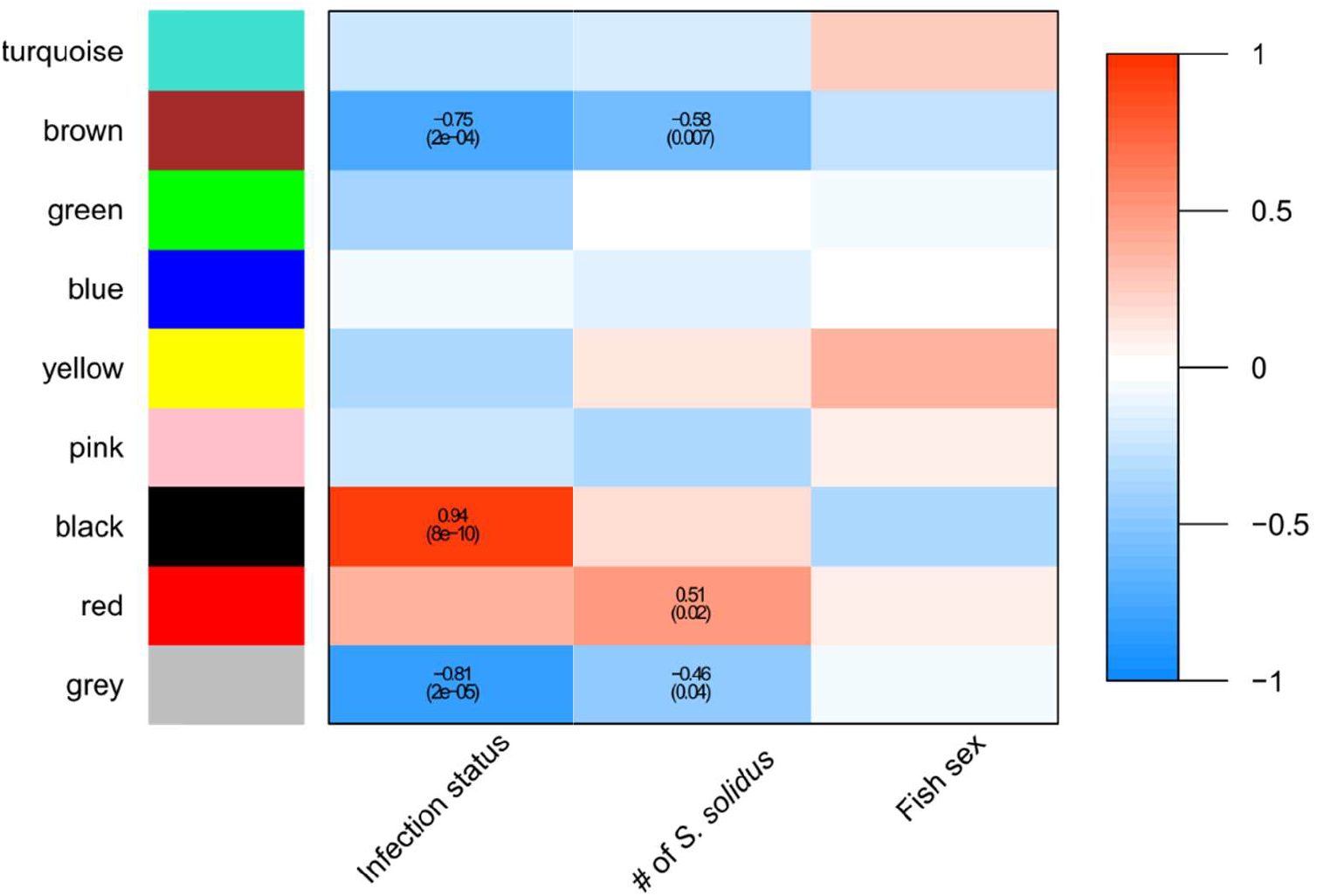
Weighed correlation gene network analysis for stickleback in Walby Lake. Only correlations with p-values less than or equal to 0.05 are presented. Cell values are Pearson Correlation coefficient plus p-values in parenthesis.

The black module was positively correlated with infection status. It was largely composed of genes associated with innate immunity (e.g., *serpinb3*, *s100a10a*, *c1q*, *anxa2a*, *f5/8 type c* DCP, *csf3r*), adaptive immunity (e.g., *myo5ab*, *f5/8 type c* DCP) and the complement system (e.g., *c1q*, *anxa2a*) (Figure 3, Table S2). One of the central genes of this module was the calcium binding gene *S100a10a*, whose protein is involved in macrophage migration and invasion (He et al., 2010). Another central gene was the cysteine protease inhibitor *serpinb3*, whose protein plays an important role in the modulation of programmed cell death in inflammatory processes (reviewed in (Vidalino et al., 2009).

**Figure 3:**
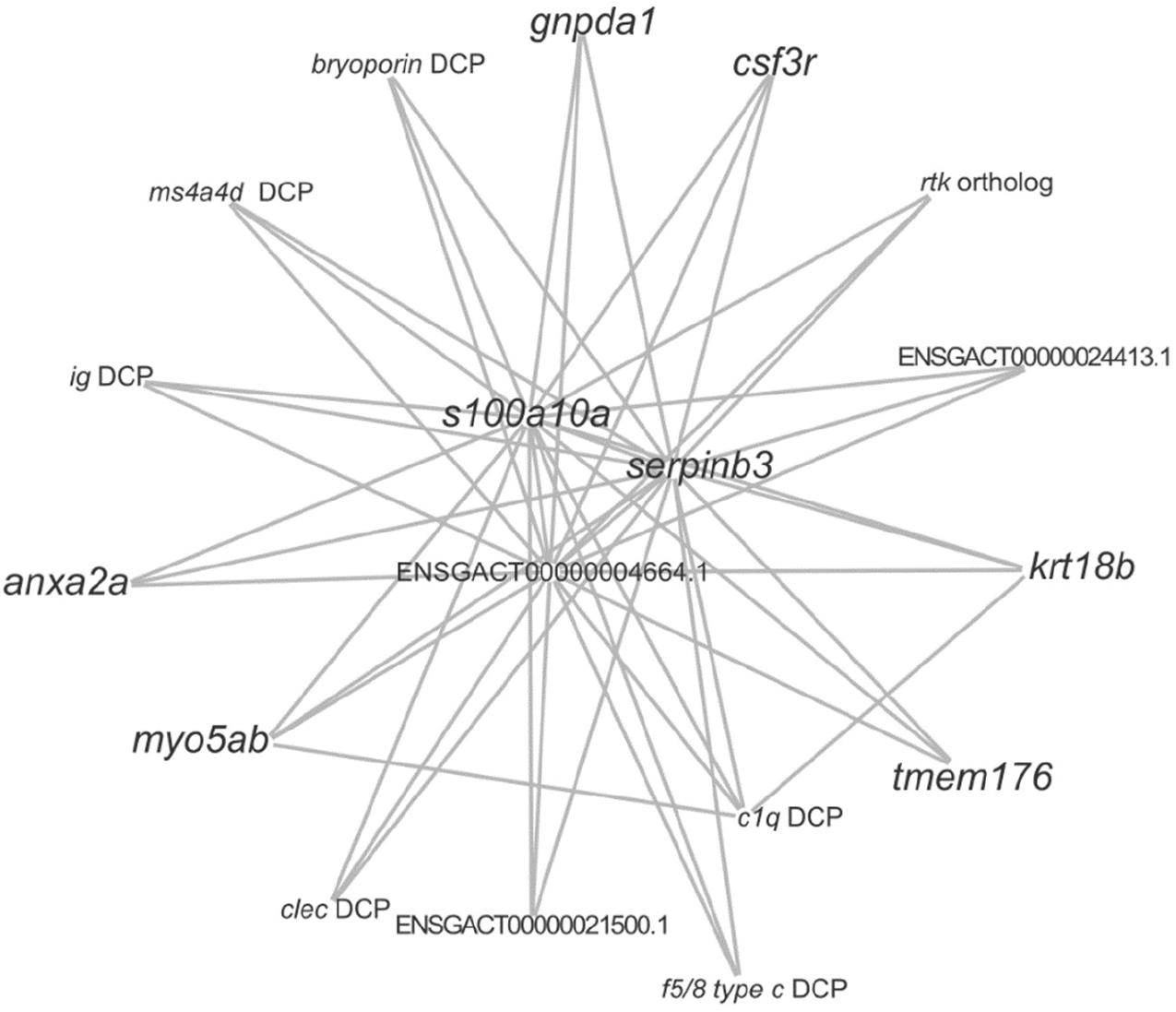
Co-expression network of genes involved in the black module for Walby Lake stickleback. Genes annotations in the larger font are confirmed via ZFIN, gene annotations in the smaller font were confirmed either via ensemble (orthologs) or UniProt (DCP). For some transcripts, no annotations were found. DCP = domain-containing protein. Gene descriptions are in the supplementary Table S2.

The brown module was negatively correlated with infection status and the number of parasites per host. Most genes in the brown module were associated with the adaptive immune response (e.g., *cd79a*, *klf2a*, *pstpip1a*, *rab29*). For instance, *rab29*, a GTPase involved in the positive regulation of T cell receptor signaling was down-regulated in infected fish. The central gene of the brown module was a class II histocompatibility antigen ortholog (*Scleropages formosus*) (ENSGACT00000000434.1, log2-fold change = -1.0459, padj = 0.001498), which plays a central role in antigen presentation to T cells (Figure 4, Table S3).

**Figure 4:**
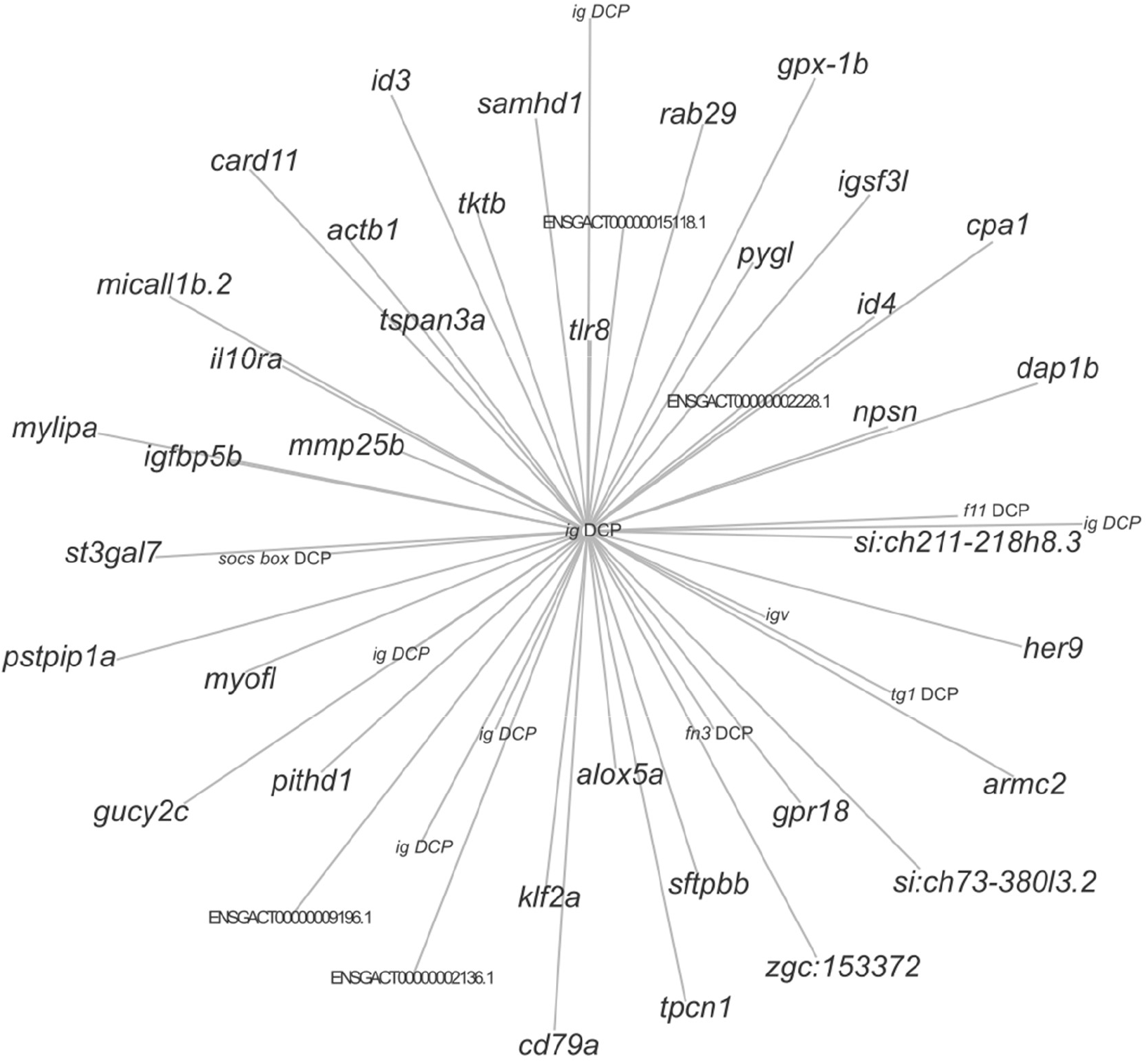
Co-expression network of genes involved in the brown module for Walby Lake stickleback. Genes annotations in the larger font are confirmed via ZFIN, gene annotations in the smaller font were confirmed either via ensemble (orthologs) or UniProt (DCP). For some transcripts, no annotations were found. DCP = domain-containing protein. Gene descriptions are in the supplementary Table S3.

The grey module was negatively correlated with both infection status and number of *S. solidus* per host. Most of the genes in the grey module were associated with a variety of components of the stickleback immune system *(e.g., c3b.2, ig, txnipa, at-like DCP, tapbp ortholog, scy DCP, hvcn1)* (Figure 5, Table S4). Other genes were associated with proteolysis *(psmb9a, psmb13a*) or calcium and iron transport/binding (e.g., *fech*, *sulf2* ortholog, *ef-hand* DCP) which, although not specific to, can play an important role in the immune response. The central gene of the grey module was *phf14*, which encodes for a histone-binding protein.

**Figure 5:**
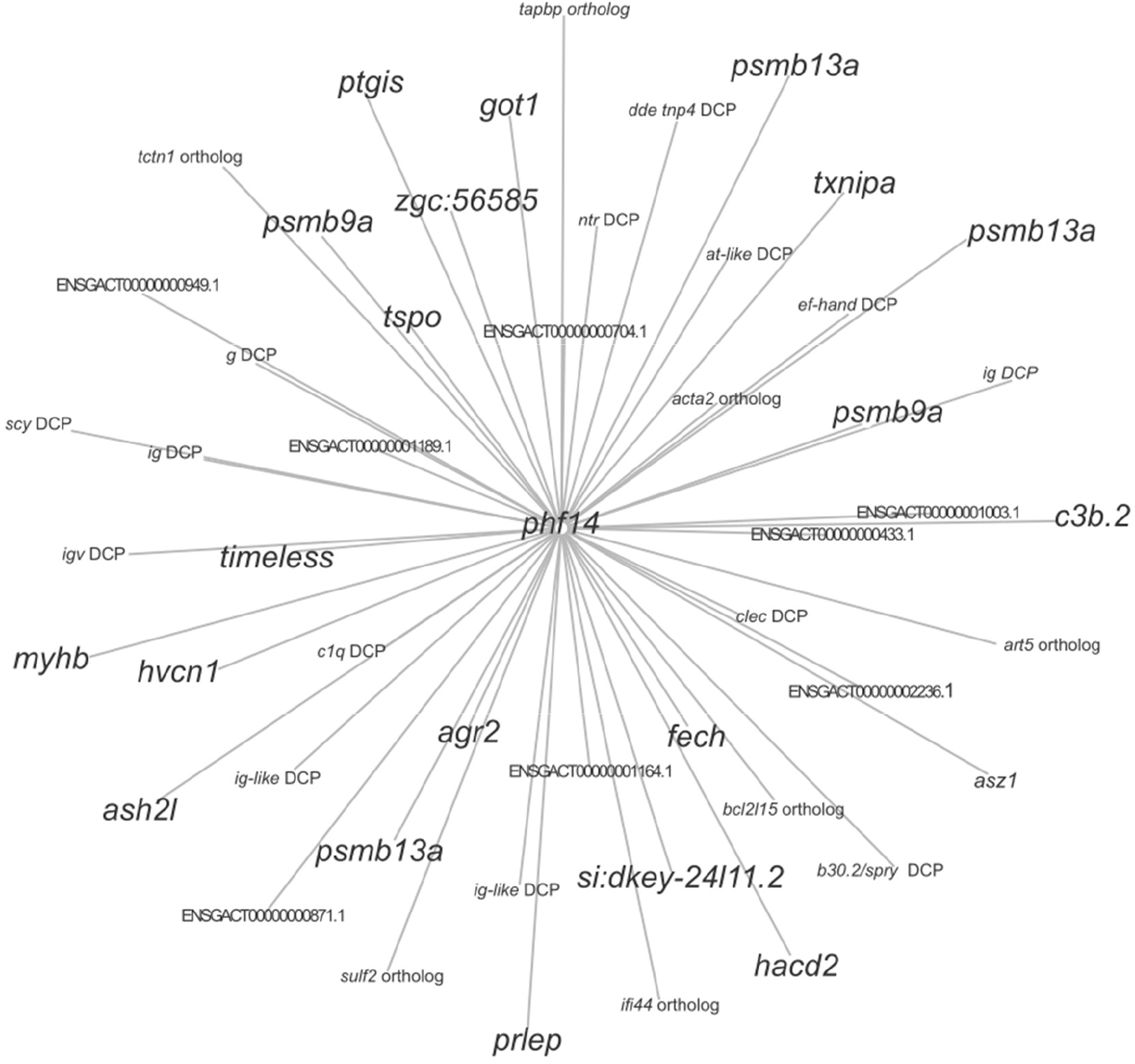
Co-expression network of genes involved in the grey module for Walby Lake stickleback. Genes annotations in the larger font are confirmed via ZFIN, gene annotations in the smaller font were confirmed either via ensemble (orthologs) or UniProt (DCP). For some transcripts, no annotations were found. DCP = domain-containing protein. Gene descriptions are in the supplementary Table S4.

Lastly, the red module was positively associated with number of *S. solidus* per host. The majority genes found in the red module were associated with interferons (e.g., *mx1*, *mxb*, *epsti1*, *cmpk2*, *hect*, *rsad*, *dhx58*, *stat1*, *mov10*, *trim21*) (Figure 6, Table S5). Furthermore, the red module also contained several genes (*ifi27l2a* ortholog, *ifi27l1*, *ifi44* ortholog, *si:ch211-197g15.10*) encoding for interferon inducible proteins, all of which are predicted to be involved in immune defense. One of the central genes in the red module encodes for an interfer-bind domain-containing protein. Interfer-bind proteins have a beta-stranded secondary structure arranged in an immunoglobulin-like beta-sandwich and can have immune-related roles, including serving as interferon receptors, tissue repair factor proteins, and interleukin receptors (Letunic et al., 2021). Another central gene in the red module encodes for protein that contains a hepn protein-coding domain, a conserved region found in the C-terminus of the chaperone *sacsin*.

**Figure 6:**
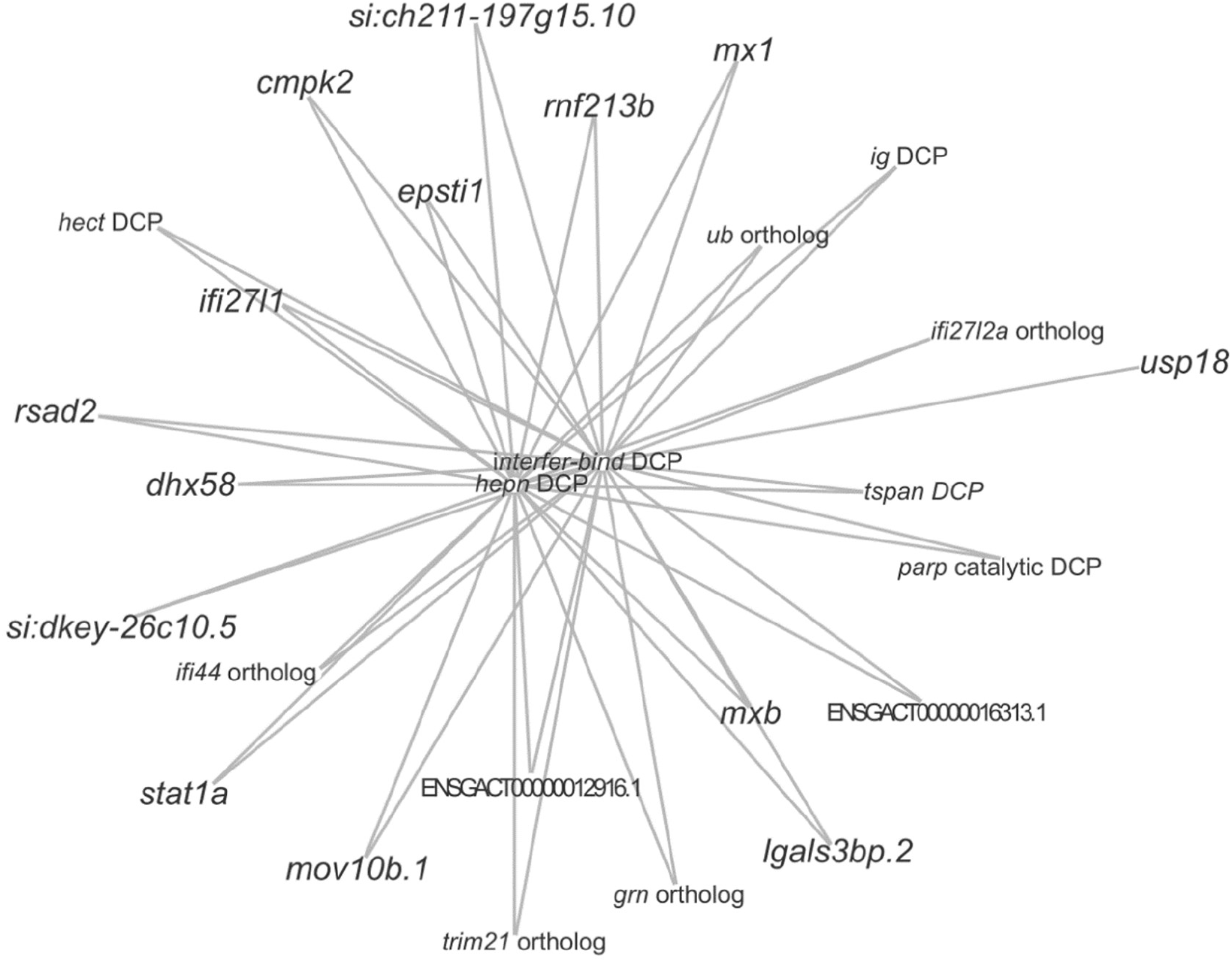
Co-expression network of genes involved in the red module for Walby Lake stickleback. Genes annotations in the larger font are confirmed via ZFIN, gene annotations in the smaller font were confirmed either via ensemble (orthologs) or UniProt (DCP). For some transcripts, no annotations were found. DCP = domain-containing protein. Gene descriptions are in the supplementary Table S5.

### Gene expression and WGCNA in Rocky Lake stickleback, a low infection population

To identify genes that contribute to parasite susceptibility in a low infection stickleback population, we compared TagSeq results between uninfected and *S. solidus*-infected Rocky Lake stickleback. Because Rocky Lake stickleback frequently encapsulate *S. solidus* into fibrotic tissue, we compared gene expression levels of uninfected fish, infected fish with non-encapsulated parasites and infected fish that had encapsulated the parasite in fibrotic tissue. TagSeq reads produced 13,310 genes for Rocky Lake samples, which is comparable to other studies of this nature (Fuess et al., 2021c; Lohman et al., 2017a). We did not find any genes to be differentially expressed between the three sampling groups except for *claudin*. *claudin* was up-regulated (ENSGACT00000005840.1, log 2-fold = 20.4025, padj = 2.02E-05) in male infected stickleback that did not encyst their parasite (Wald, p < 0.1 after 10% FDR correction); it codes for a transmembrane protein found in tight junctions.

As with Walby Lake stickleback, we employed network co-expression analysis to identify correlation patterns among gene sets, as well as between gene sets and sample features (e.g., infection status, stickleback sex, co-infection by other parasites). We identified 13 co-expression modules, four of which (pink, black, tan, grey) were associated with stickleback sex (Figure 7). The presence of encysted *S. solidus* and presence of peritoneal fibrosis were both negatively associated with the green module. The number of non-encysted (free) *S. solidus* was negatively associated with the blue module and lastly, the number of encysted *S. solidus* was negatively associated with the grey module. No module was associated with the presence of other parasites in the sampled fish or the presence of non-encysted *S. solidus*.

**Figure 7:**
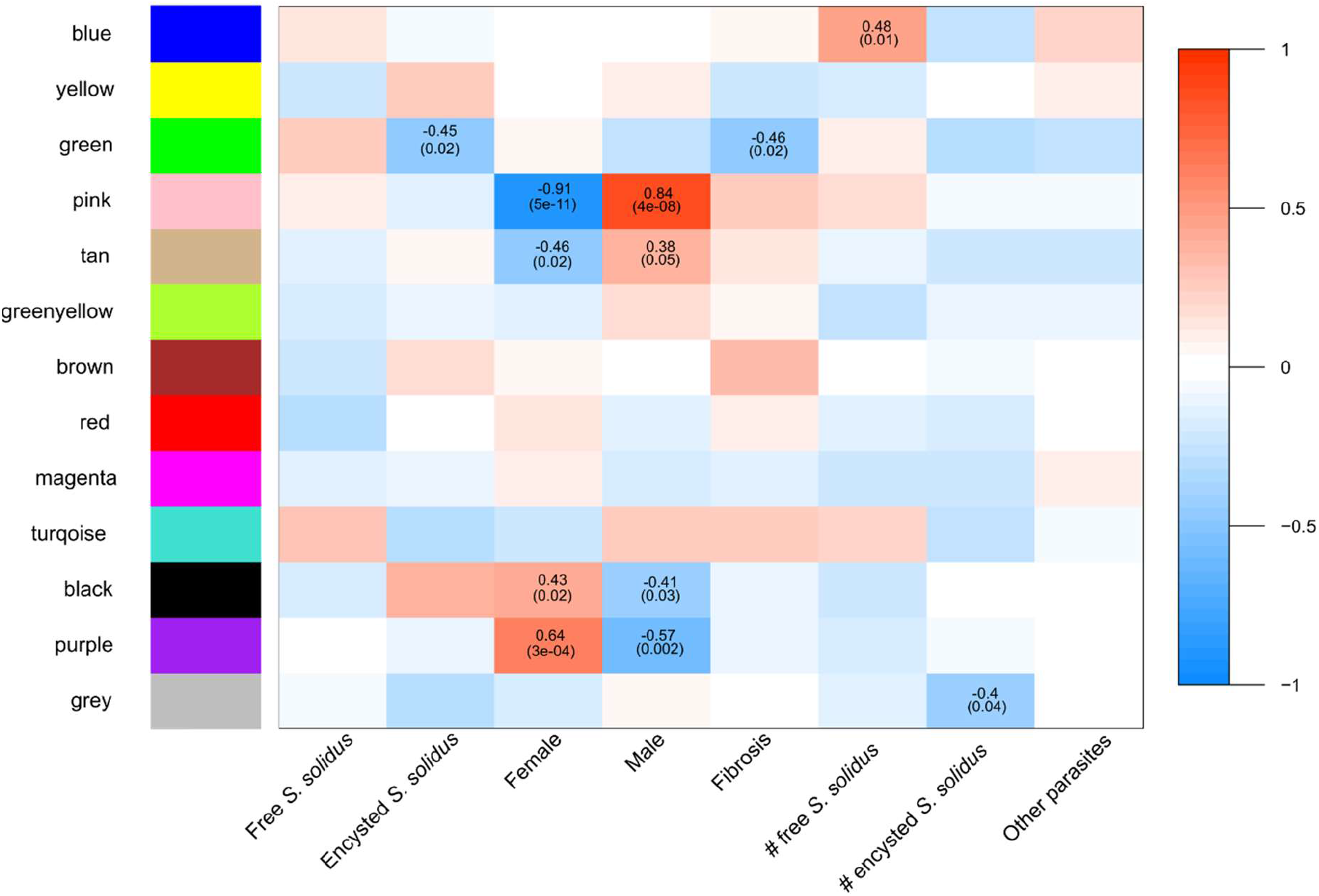
Weighed correlation gene network analysis for stickleback in Rocky Lake. Only correlations with p-values less than or equal to 0.05 are presented. Cell values are Pearson Correlation coefficient plus p-values in parenthesis.

The green module was negatively associated with the presence of encysted *S. solidus* as well as the general presence of peritoneal fibrosis. Fittingly, several genes in the green module were related to fibrosis formation (e.g., *kank4*, *fn3* DCP, *col1a1*, *cers2b*, *gpnmb*). *col1a1* (*collagen, type I, alpha 1a*) for example is currently under debate as a marker for cardiac fibrosis in humans (Hua et al., 2020), and research has shown that the inhibition of Ceramide (*cers2b*, *ceramide synthase 2b)* accumulation improves fibrosis in liver tissue samples in rats (Jiang et al., 2019). *gpnmb* (*glycoprotein (transmembrane) nmb*) is associated with scleroderma fibrosis (Palisoc et al., 2022). Other genes were associated with heme synthesis (e.g., *uros*, *ppox*, *cpox*), iron or calcium transport (e.g., *cracr2b*, *slc4a1a*, *fer DCP*) and the immune system (e.g., *bcl6*, *c1q*, *bpi* DCP, *tspo*, *gstm4*). The central gene in the green module was a class II histocompatibility antigen ortholog (ENSGACT00000000198.1, *Poecilia formosa*) (Figure 8, Table S6), which plays a central role in antigen presentation. Several of the genes involved in the green module in Rocky Lake fish were also found in the grey module in Walby Lake fish (*c1q* DCP, *ig* DCP, *tspo*).

**Figure 8:**
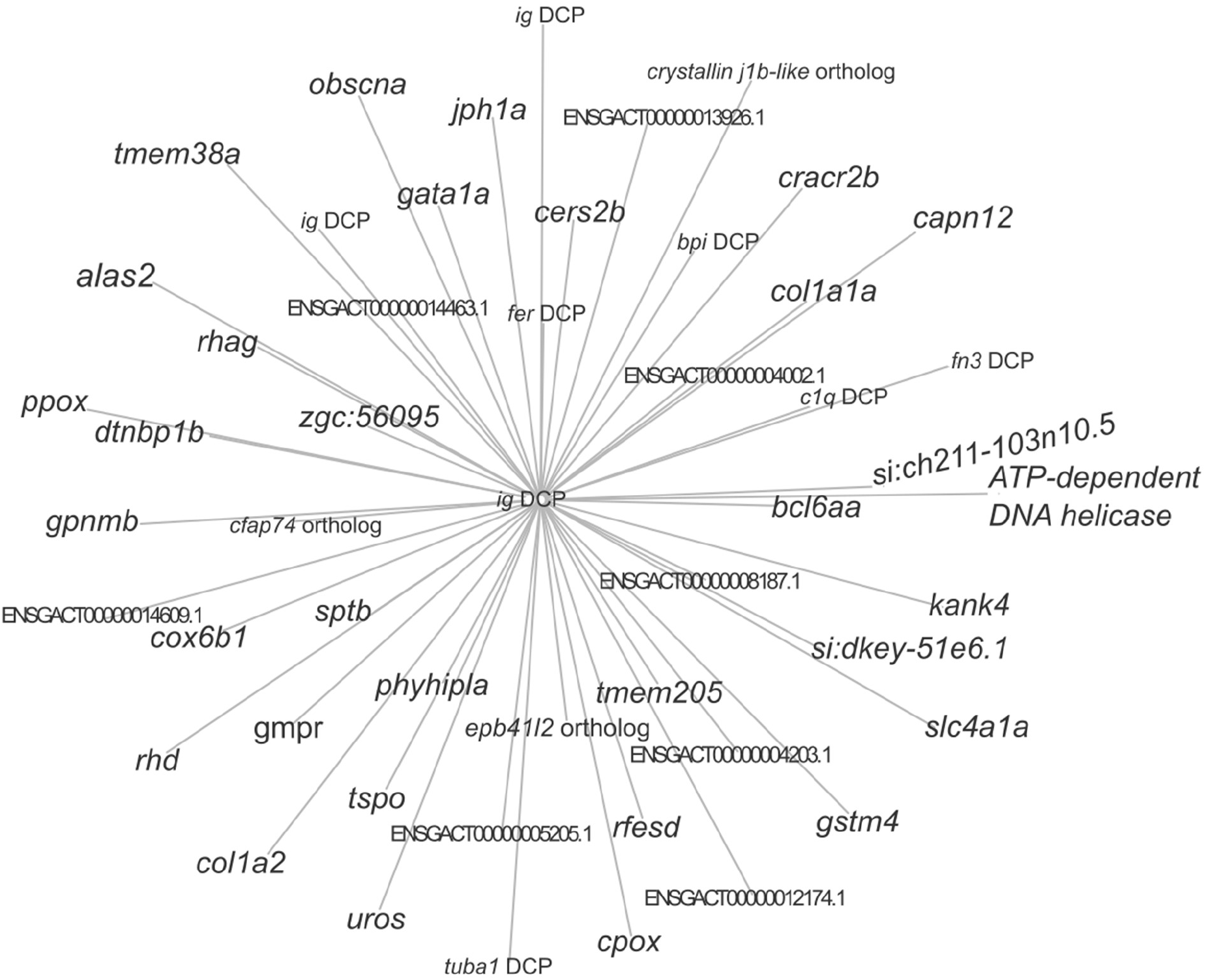
Co-expression network of genes involved in the green module for Rocky Lake stickleback. Genes annotations in the larger font are confirmed via ZFIN, gene annotations in the smaller font were confirmed either via ensemble (orthologs) or UniProt (DCP). For some transcripts, no annotations were found. DCP = domain-containing protein. Gene descriptions are in the supplementary Table S6.

The grey module was negatively associated with the number of encysted *S. solidus* per stickleback host. Many genes in the grey module lacked annotation (Figure 9, Table S7), while other genes were associated with the modulation of the stress response (*scgn*). Two of the more central genes in this module were associated with G protein receptor signaling (*acp6*, *rgs13*).

**Figure 9:**
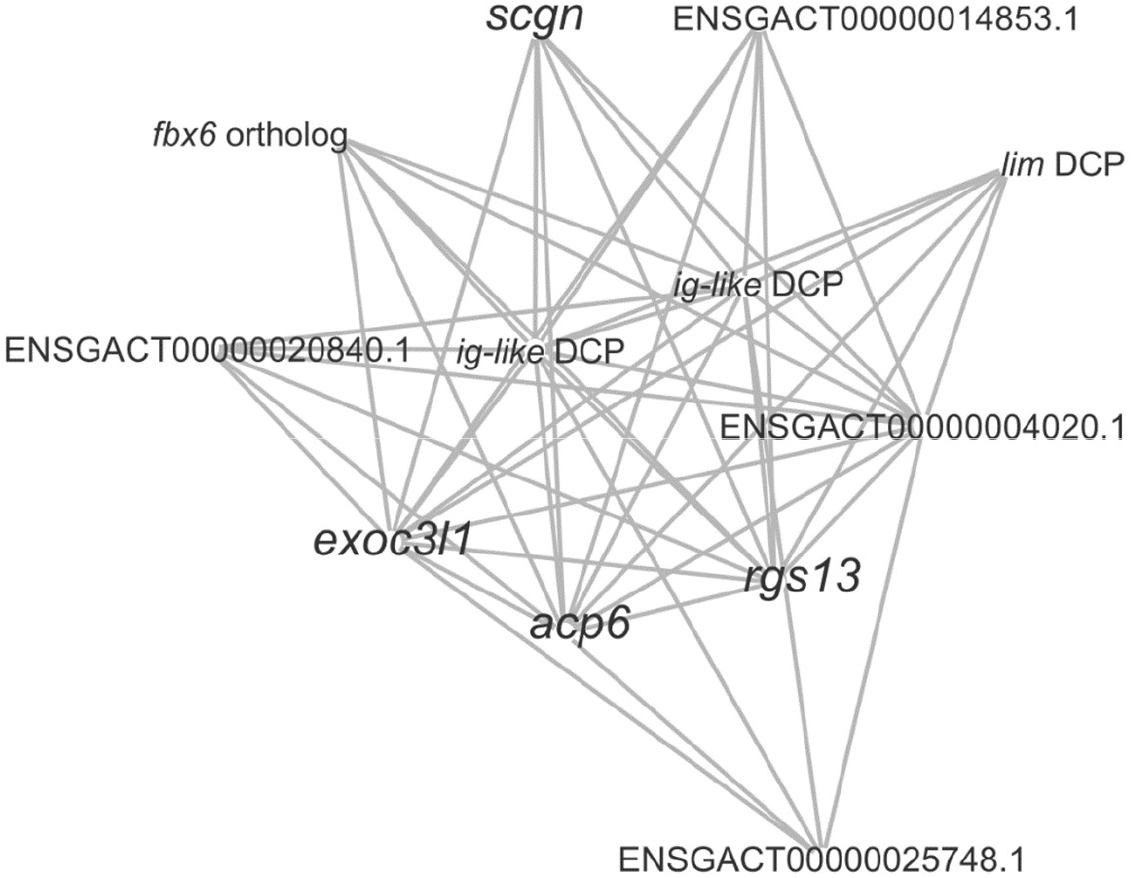
Co-expression network of genes involved in the grey module for Rocky Lake stickleback. Genes annotations in the larger font are confirmed via ZFIN, gene annotations in the smaller font were confirmed either via ensemble (orthologs) or UniProt (DCP). For some transcripts, no annotations were found. DCP = domain-containing protein. Gene descriptions are in the supplementary Table S7.

The blue module was negatively associated with the number of non-encysted *S. solidus* per host. Several of the genes in this module are involved in protein transport and translocation (e.g., *ssr1*, *cuta*, *nedd9*, *abcc6b*). Other genes are involved in innate (e.g., *il-8* DCP, *commd3*, *clec*, *timp2a*, *cxcr1*, *ita4h*, *npsn*) and adaptive (*mhc I-like ar-recog* DCP, *clec*, *chia*) immunity as well as gene expression and the cell cycle (e.g., *med11*, *sympk*, *srsf2*, *mad2l1*, *nedd9*, *rps20*). The central gene in the blue module was a mitochondrial intermembrane chaperone (*timm8b*, *translocase of inner mitochondrial membrane 8 homolog B*) that is involved in the import of multi-pass transmembrane proteins (Figure 10, Table S8). Several genes found in the blue module in Rocky Lake stickleback were also present in either the grey (*psmb13a*, *clec* DCP, *fech*) or the brown module (*ig-like* DCP, *npsn*) of Walby Lake stickleback.

**Figure 10:**
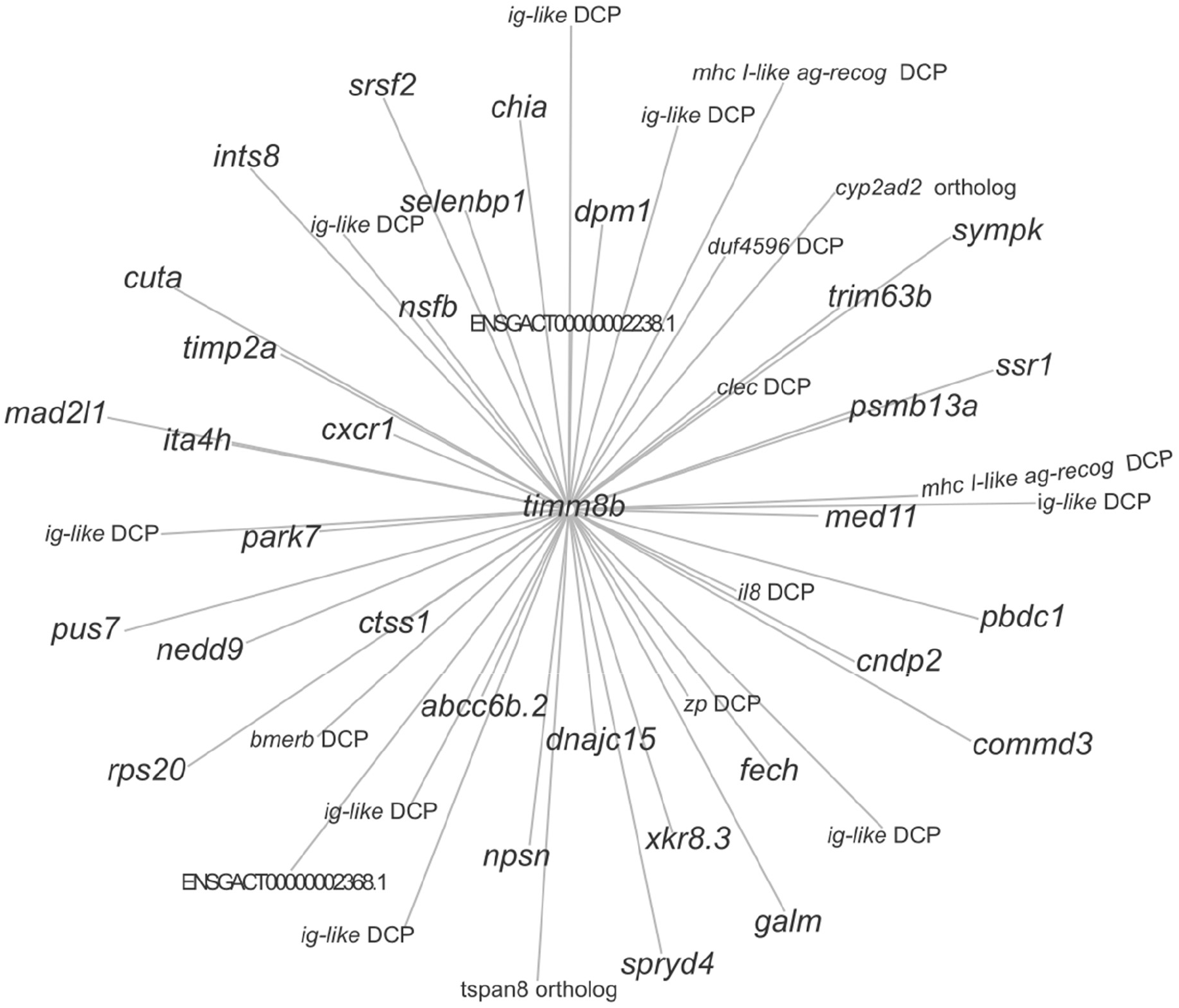
Co-expression network of genes involved in the blue module for Rocky Lake stickleback. Genes annotations in the larger font are confirmed via ZFIN, gene annotations in the smaller font were confirmed either via ensemble (orthologs) or UniProt (DCP). For some transcripts, no annotations were found. DCP = domain-containing protein. Gene descriptions are in the supplementary Table S8.

### Comparing gene expression in uninfected stickleback in Rocky Lake and Walby Lake

To assess differences in baseline immune gene expression between a high and a low infection population, we compared gene expression between uninfected fish in Rocky Lake and Walby Lake. TagSeq reads produced 12,526 genes for uninfected fish in Rocky Lake and Walby Lake. Of these 12,526 genes, 10.15% were up-regulated and 9.92% were down-regulated in Rocky Lake fish compared to Walby Lake fish (Wald, p < 0.1 after 10% FDR correction). We found many genes associated with innate immunity, and specifically with inflammation, to be more highly expressed in the uninfected Rocky Lake fish (high infection population) than in Walby Lake fish (Table 2, Table 4). For example, *galectin* is a family of lectins that act as pattern recognition receptors detecting conserved pathogen-associated molecular patterns of various parasites, helping the host to mediate parasite recognition and activation of immune responses, such as apoptosis of host cells (reviewed in (Shi et al., 2018). Another example is *alox5ag* (*arachidonate 5-lipoxygenase activating protein*), which is required for leukotriene synthesis; Leukotrienes are inflammatory mediators produced in leukocytes. Genes involved in other parts of the immune system were also up-regulated in Rocky Lake fish, for example *fhl-1* (*complement factor h-like*) which is involved in the complement pathway and protects host surfaces from uncontrolled complement attack.

**Table 4:** Selection of top differentially expressed genes associated with immune functions in uninfected Rocky Lake stickleback compared to uninfected Walby Lake stickleback. The log 2-fold change refers to the differences in uninfected Rocky Lake fish compared to uninfected Walby Lake fish. The p-value (padj) is adjusted for multiple testing via Benjamini-Hochberg (10% FDR).

| Ensembl Transcript ID | Gene name | Un – Log2-fold change | Un – *padj* | Putative Immune function |
| --- | --- | --- | --- | --- |
| ENSGACT00000008899.1 | *interferon alpha-inducible protein 27-like protein 2a (ifi27a)* | 4.9706 | 2.49E-05 | Innate Immunity; Apoptosis |
| ENSGACT00000016708.1 | *galectin* | 4.0652 | 1.94E-75 | Innate Immunity & Adaptive Immunity; Immune homeostasis |
| ENSGACT00000008904.1 | *interferon alpha-inducible protein 27 like 1 (ifi27l1)* | 3.6968 | 4.90E-05 | Innate Immunity; Apoptosis |
| ENSGACT00000004261.1 | *complement factor h-like (fhl-1)* | 3.6022 | 4.17E-08 | Complement System |
| ENSGACT00000007295.1 | *serpinb1* | 3.2183 | 1.28E-08 | Innate Immunity; Inflammation |
| ENSGACT00000004464.1 | *s100 calcium-binding protein a10 (s100a10a)* | 1.9039 | 2.32E-09 | Innate Immunity; Inflammation & TLR Signaling |
| ENSGACT00000018254.1 | *granulocyte colony stimulating factor 3 (csf3r)* | 1.6075 | 1.95E-11 | Innate Immunity; Granulocyte Maturation |
| ENSGACT00000027217.1 | *arachidonate 5-lipoxygenase activating protein (alox5ap)* | 0.9382 | 4.61E-08 | Innate Immunity; Inflammation |
| ENSGACT00000020376.1 | *chemokine like factor (cklf)* | 1.0324 | 2.10E-09 | Innate Immunity; Inflammation |

## Discussion

The goal of this project was to identify mechanisms underlying differences in *S. solidus* susceptibility in different stickleback populations with a focus on immune gene expression. We characterized head kidney gene expression in one high infection (Walby Lake) and one low infection (Rocky Lake) population as well as the differences between the two populations.

### Differences in immune gene expression in Walby Lake stickleback

In infected Walby Lake stickleback, we observed the up-and down-regulation of many genes involved in host immunity and parasite defense (Table 2, Table 3). By choosing to collect data from naturally-infected populations, we had no control over infection timing and were not able to determine how long individual fish had been infected. Previous work has shown that different parts of the stickleback immune system are up- or down-regulated at various time points after the encounter with *S. solidus* (Scharsack et al., 2004b, 2007; Wohlleben et al., 2018). Even though we did not weigh each *S. solidus* individually, most appeared to be fully infective for the final host (Tierney and Crompton, 1992) and likely several weeks in age. This makes it the more surprising that we found several genes associated with innate immunity (e.g, *fn*, *cpvl*, *fxyd5*, *csf3r*, *anxa2a*) to be up-regulated as macroparasites should be too large to be cleared by the innate immune system at this point (Scharsack et al., 2007; Secombes and Chappell, 1996). However, although the innate immune system is the first line of defense in many animals, there is some evidence that the innate immune system may be activated late (after several weeks) during infection with *S. solidus* in stickleback (Haase et al., 2016; Piecyk et al., 2019; Scharsack et al., 2007). Early during infection, *S. solidus* can evade immune detection or actively down-regulate stickleback immune activation (Wedekind and Little, 2004). Furthermore, the proliferation of head kidney monocytes, precursors of granulocytes and macrophages is only mobilized seven days post infection (Scharsack et al. 2007). Accordingly, we found a key component of innate immunity, the *granulocyte colony stimulating factor 3* (*csf3r*) to be up-regulated in our samples. CSF3R plays a vital role in differentiation and proliferation of granulocytes; its activation and mobilization is a typical response in fish exposed to helminth parasites (Balla et al., 2010; Nie and Hoole, 2000; Reite and Evensen, 2006; Scharsack et al., 2004b; Sharp et al., 1992). Although granulocyte mobilization may be effective in the early stages of *S. solidus* infection, once the parasite reaches a certain size, granulocytes cannot clear the infection (Scharsack et al., 2004). *csf3r* is also part of the black Walby module, which we identified via weighted gene co-expression analysis. Other genes in the black Walby module are positively associated with inflammation (e.g.: *serpinb3*, *s100a10a*, *c1q*, *anxa2a*) and the complement system (e.g.: *c1q*, *anxa2a*) (Figure 3). The complement system plays an important role in inflammation and the killing of pathogens and parasites through the organization and activation of phagocytosis. Genes of the complement cascade seems to generally be up-regulated at later stages of *S. solidus* infections (Haase et al., 2016; Piecyk et al., 2019). One of the central activators of the complement cascade is the *complement component 1q* (*c1q*), which we found to be up-regulated in infected Walby stickleback. The up-regulation of *c1q* points towards a general activation of the complement system. On top of being central to the activation of the complement system, *c1q* is also involved in the modulation of the T cell response, bridging innate and adaptive immunity (Bennett et al., 2017; Nakao et al., 2011; Sontheimer et al., 2005). Overall, it seems that the cellular innate immune response and the complement cascade are up-regulated late during in the infection with *S. solidus* in Walby Lake fish. This might suggest a strategy by the parasite to delay the innate immune response until it cannot be cleared anymore.

A previous study in European stickleback populations found that monocyte proliferation was down-regulated in infected stickleback, indicating a suppression of the stickleback adaptive response by *S. solidus* (Scharsack et al., 2007). Similarly, we observed a down-regulation of genes involved in T cell receptor signaling (e.g.: *lck*, *zap-70*, *dgkab*, *irf4a*, *skap1, tcr-β*) in infected Walby Lake stickleback. T cells (Tc, Th1, Th2, Th17, Treg) are important players in the adaptive immune response. The T cell receptor recognizes peptides presented by the MHCs and is therefore essential for an effective host response to parasites and pathogens (Lo et al., 2018; van der Merwe and Dushek, 2011). The suppression and even down-regulation of lymphocytes and specifically T cells is a common trend in *S. solidus* infections (Fuess et al., 2021; Scharsack et al., 2007). Although helminth infections typically induce a Th2 response, gene expression data from the head kidneys of stickleback infected with *S. solidus* does not point towards this typical Th2 response (Orr et al., 1969; Scharsack et al., 2007). In the current study, expression levels of *stat4* and *stat6*, transcriptional activators needed for the Th1 and Th2 pathways, respectively, showed no differences, consistent with earlier studies (Orr et al., 1969; Scharsack et al., 2007). It is important to note that we only analyzed head kidneys to assess immune gene expression. While the head kidney is a primary immune organ in teleost fish, immune cell types are not equally expressed in every immune organ. For example, head kidneys are the primary B cell organ, but the thymus is the primary T cell organ. Only analyzing one immune organ is not enough to get a full picture of the immune response and future research should consider analyzing several tissues simultaneously.

The MHC plays a central role in presenting antigens to the adaptive immune system, and within the MHC I, proteasomes are essential for antigen presentation. We found a negative association between proteasome expression (grey Walby module) and infection status, as well as the number of parasites per host, suggesting reduced activation of the adaptive immune response. Besides their role in antigen presentation (adaptive immunity), proteasomes are also involved in the regulation of the innate immune response. Upon stimulation by pro-inflammatory cytokines, immunoprotoeasomes, formed by the exchange of catalytic sited in the standard 20S proteasome core complex, participate in degrading intracellular proteins for presentation by MHC class I molecules (reviewed in Kammerl & Meiners, 2016). We found down-regulation of proteasomes as well as a class II histocompatibility antigen ortholog (Figure 4). The MHC class II protein complex has been linked to sticklebacksusceptibility to *S. solidus* infections (Haase et al., 2016; Kurtz et al., 2004; Lohman et al., 2017; Wegner et al., 2006); in German stickleback, high parasite load was positively correlated with MHC II expression (Kurtz et al., 2004). In our study, we could not differentiate between MHC haplotypes due to the nature of TaqSeq analysis, which generates single fragment from the 3’ end of each transcript (= tag) and quantifies gene expression based on tag abundance. The expression levels of the immunoglobulin superfamily member *cd83*, and promotes the expression of MHC II (Grosche et al., 2020; Tze et al., 2011), did not differ in between infected and uninfected stickleback in the high infection Walby Lake population. Overall, the downregulation of MHC II antigen and proteasome expression might impair the antigen-presenting capacity of antigen-presenting cells, which may allow *S. solidus* to evade the host’s adaptive immune response.

We found several genes to be correlated with both infection status and the number of *S. solidus* per host. Stickleback analyzed in this experiment harbored between 1 and 44 parasites, with most hosts harboring two or three. Multiple infections by *S. solidus* larvae in a single stickleback are common in nature. On the one hand, co-infecting parasites compete for the same host resources, possibly limiting parasite fitness. On the other hand, a simultaneous attack by several parasites may pose an additional challenge to the host immune system, potentially increasing parasite fitness (Karvonen et al., 2012). As of now, we cannot predict how intra-host competition affects the stickleback-*S. solidus* interaction, and previous studies have found mixed results. For example, work on Norwegian stickleback populations showed that stickleback with multiple *S. solidus* infections had higher relative body conditions, suggesting that solo infections are more harmful than multiple infections (Nordeide and Matos, 2016). Conversely, host body condition decreases significantly with increasing number of parasites per host in Walby Lake fish (Heins et al., 2019). Our study supports this finding, as immune genes are increasingly down-regulated with an increasing number of parasites per host, indicating a progressively negative impact of infection burden on Walby Lake stickleback.

We saw a positive correlation between *interferon* (IFN) expression, cytokines typically produced in response to intracellular infections, including viruses and can interfere with viral replication, and number of parasites per host. Intrestignly, previous studies in mice infected with the intestinal nematodes (*Heligmosomoides polygyrus* and *Nippostrongylus brasiliensis*) type I interferon and interferon-stimulated genes were up-regulated, leading to enhances parasite fecundity and delayed parasite clearing (McFarlane et al., 2017; Urban et al., 1993). Interestingly, blocking of IFN signaling reversed these effects (Urban et al., 1993). The up-regulated expression and/or activity of type I interferons might represent a strategy employed by helminth to avoid the host immune response, benefiting the parasite (reviewed in (Silva-Barrios and Stäger, 2017). While it might seem surprising, previous research has shown that some parasites can stimulate parts of the host immune system to counteract specific responses (reviewed in (Chulanetra and Chaicumpa, 2021). Something similar may have occurred in our study, where fish with more parasites had a higher expression of IFN genes, possibly diluting a targeted response against *S. solidus*.

The GO term analysis for Walby Lake fish (Figure 1) showed a slight change in immune gene-related activity following infection. This could be an artifact of missing annotations when the GO terms were created. Many of our highly differentially expressed genes are associated with the immune response although they are not associated with immune gene GO terms. For this reason, we focused on candidate genes and analyzed the top 100 differentially expressed genes. This finding highlights the importance of exploring individual gene expression patterns as GO term analyses alone may not capture all relevant genes, especially in non-model organisms. GO terms are a great tool for analyzing large -omic datasets, but they can introduce biases when our knowledge underlying the GO terms is incomplete (Gaudet and Dessimoz, 2017).

### Immune gene expression levels in Rocky Lake stickleback

In contrast to Walby Lake stickleback, we only found one gene to be differentially expressed between infection statuses in Rocky Lake stickleback. *Claudin*, which was up-regulated in male infected stickleback, codes for a tight junction protein, which are proposed to be effectors and targets of mucosal immune regulation (Zuo et al., 2020). Tight junctions play an essential role in the intestinal barrier function, regulating the paracellular movement of various substances (e.g.: ions, solutes, water). They also act as a first line of host defense against pathogen/parasite intrusion from the intestinal lumen to the underlying tissue. As *S. solidus* has to penetrate the gastrointestinal mucosal wall to establish itself in the body cavity, this could explain the up-regulation.

Rocky Lake stickleback are rarely infected with *S. solidus*, which made it difficult for us to balance sampled stickleback for confounding factors such as stickleback sex or co-infections by other parasites. In fact, when we analyzed the dataset via WGCNA we found considerable impact of stickleback sex on Rocky gene co-expression; four (pink, black, tan, purple) of the 13 identified co-expression modules are associated with stickleback sex, while only one (green) is with infection status (Figure 7). Sex-biased parasitism has been reported in the past (Bolnick et al., 2020; Natsopoulou et al., 2012; Reimchen and Nosil, 2001), but its presence varies among stickleback populations, with either males or females being more heavily parasitized (Natsopoulou et al., 2012) (Reimchen & Nosil, 2001). When organisms experience different infection pressure, they can evolve different counter mechanisms, which is why we might see many gene co-expression networks that differ between stickleback sexes. In the future it will be important to determine sex-infection interactions by collecting data on the relative numbers of female and male infected fish in both sampled lakes.

Rocky Lake stickleback regularly induce peritoneal fibrosis upon parasite exposure, and we found encysted *S. solidus* in the body cavity. Fibrosis formation as defense against *S. solidus* has developed in several stickleback freshwater populations (Fuess et al., 2021c; Hund et al., 2022; Lohman et al., 2017a; Vrtílek and Bolnick, 2021b; Weber et al., 2022), inhibiting cestode growth and potentially killing the parasite. We identified several fibrosis-associated genes (green Rocky module) that were negatively associated with the presence of encysted *S. solidus* (Figure 8). Interestingly, one of these genes codes for a Fibronectin domain-containing protein (ENSGACT00000008537.1), which we also found in the brown module in Walby Lake (negatively associated with infection status and number of *S. solidus* per host). This contrasts with previous findings of up-regulated fibronectin in infected fish from British Columbia (Fuess et al., 2021). Fibronectin, an extracellular matrix protein, has been linked to increasedfibrosis (Valiente-Alandi et al., 2018). The negative association of the green Rocky module in our study suggests an ongoing effort of the fish to reduce peritoneal fibrosis, as it is a costly defense mechanism that reduces fish fecundity and overall fitness.

Several genes that are predicted to be involved in the MHC I immune defense (*clec*, *commd3*, *mhc I-like ar-recog* 2x) were negatively associated with the number of non-encysted *S. solidus* per host (blue Rocky module). This aligns with the reduced expression of adaptive immune response genes observed in infected Walby Lake stickleback. The blue Rocky module also contained innate immune system genes such as *chia* (*chitinase acidic*), which plays an important role in the Th2 immune response. It is important to note, however, that genes were not down-regulated in Rocky Lake stickleback (as defined in the DEseq2 package) but rather part of a gene co-expression network that was negatively associated with infection. Still, this might indicate the active manipulation of the adaptive immune response by the parasite.

Analyzing naturally-infected stickleback resulted in us collecting fish at various points post *S. solidus* exposure. Rocky Lake is considered a medium to low infection population, and infection rates were found to vary between 2% and 10% (Wohlleben, 2022) over an 11-year sampling period. This made it difficult to collect stickleback that harbored *S. solidus* of similar size and developmental stage for the present study. Previous research has shown that different parts of the stickleback immune system are up- or down-regulated at different time points after the exposure to *S. solidus* (Piecyk et al., 2019; Scharsack et al., 2007; Wohlleben et al., 2018). It is therefore likely that variation introduced by co-founding variables introduced noise, which made it difficult for us to detect differences between infection groups. Furthermore, Rocky Lake fish might show a higher level of plasticity in their base immune gene expression levels due to factors in their environment that are currently unknown to us. Laboratory studies control for (a)biotic factors to increase the likelihood of detecting infection-induced expressen differences. However, studying immunity in natural environments is crucial to understand the immune system’s evolution and functioning. Previous work found that stickleback that were transplanted from their home lake to new environments showed an immune gene expression resembling that typical of individuals in the destination lakes, underpinning the importance of studying immunity in the natural environment where the immune system has evolved and is actively functioning. While we ran the risk of overlooking differentially expressed immune genes because of environmental variation, we wanted to avoid measuring atypical immune traits, induced by keeping stickleback in laboratory environments. Fruther research on Rocky Lake stickleback, including the monitoring of *S. solidus* growth rates and conducting repeat experments while tracking parasite size for aproxy of infection timing, is important to advance our understanding of this particular host population.

### Gene expression between populations

The stickleback populations in the present study were chosen because of their difference in *S. solidus* susceptibility: Walby Lake stickleback typically show high infection rates, while Rocky Lake stickleback typically show low infection rates (Heins et al., 2010, 1999; Wohlleben, 2022). Therefore, we were not only interested in gene expression within population between infection statuses but also between populations. To assess baseline gene expression, we compared TagSeq results in uninfected fish. We found that about 20% of genes were differentially expressed between populations, 10.15% of which were up-regulated and 9.92% were down-regulated in Rocky Lake fish compared to Walby Lake fish (Figure 11). Interestingly, many genes that we found up-regulated in Rocky Lake fish were associated with the innate immune response and specifically with inflammation (Table 2, Table 3, Table 4). In addition to the high expression of innate immune genes (e.g., *csf3r*, *c1q*, *alox5ap*, *cxcl19*), we also identified genes that protect against immunopathology. For example, *serpinb1* is a neutrophil protease inhibitor that primarily acts to protect host cells from proteases released in the cytoplasm during stress or infection (Choi et al., 2019). In Rocky Lake stickleback, we may expect pathways required to prevent immunopathology to be up-regulated if their innate immune system is more activated. As the innate immune response is the first line of defense against pathogens, this relatively high expression of innate immune genes might indicate a higher risk of parasite or pathogen exposure in Rocky Lake. We unfortunately do not have ecological data on parasite communities in the two lakes but know from our own experience that Rocky Lake stickleback are commonly infected with black spot parasites, which is rarely observed in Walby Lake.

### Conclusions and future outlook

We observed between-population variation in response to *S. solidus* infection in naturally-infected stickleback from Walby Lake (high infection) and Rocky lake (low infection). Walby Lake stickleback exhibited differential gene expression between infection statuses, with up-regulation of genes related to the cellular innate immune response and complement cascade, and down-regulation of genes associated with adaptive immunity, suggesting immune manipulation by the parasite. Immune gene expression was negatively correlated with parasite burden (number of parasites), supporting previous work indicating that a higher number of parasites has an increasing negative effect on the host.

In contrast, Rocky Lake stickleback showed high levels of peritoneal fibrosis and several fish had encysted *S. solidus*. Although only one gene showed differential expression between infection statuses, we identified changes in gene co-expression networks. Most were associated with stickleback sex, which could indicate sex-biased infection pressure or reflect an overall difference between the two sexes. The lack of differentially expressed genes may suggest higher gene expression variability in Rocky Lake fish. Interestingly, when we compared immune gene expression of uninfected fish, Rocky Lake fish showed an up-regulation of innate immune genes compared to Walby Lake fish. Future studies should investigate overall parasite communities, infection rates, and potential sex-infection interactions to better understand those differences.

## Data

Data will be made available upon acceptance in a peer reviewed journal.

## Acknowledgments

We would like to thank Susan Foster and John Baker for their input during the early stages of the project. Their expertise and advice significantly contributed to the development and direction of our research. Special thanks are due to Dr. Andrea Roth-Monzón, Dr. Jesse Weber, and Kevin neumann for their invaluable assitstance in collectin the stickleback samples in Alaska. Their dedication and support in the field were instrumental in the success of our data collection. We are grateful to Dr. Kaitlyn Mathis and Dr. Robert Drewell for their instightfull feedback throughout all stages of this work. Lastly, we would like to acknowledge the the Nipmuc people (MA) and Dena’ina Elnana and Dënéndeh tribes (AK) on whose ancestral land we conducted our research. We acknowledge their connection to the lands and express our respect for their cultures, traditions, and knowledge.

## Author Contributions

AMW, NPM, and NCS collectively conceived and designed the study. AMW took the lead in data gathering and conducted most of the statistical analysis. JPT provided valuable assistance in the bioniformatics analysis. AMW, NPM and NCS wrote the article.

## Ethical Standards

The authors assert that all fish collections were made in accordance with IACUC approved animal use protocols from Clark University (013R) and approved Aquatic Resource permits from the Alaska Department of Fish and Game (SF2021-106).

## Appendix

**Table S1:**
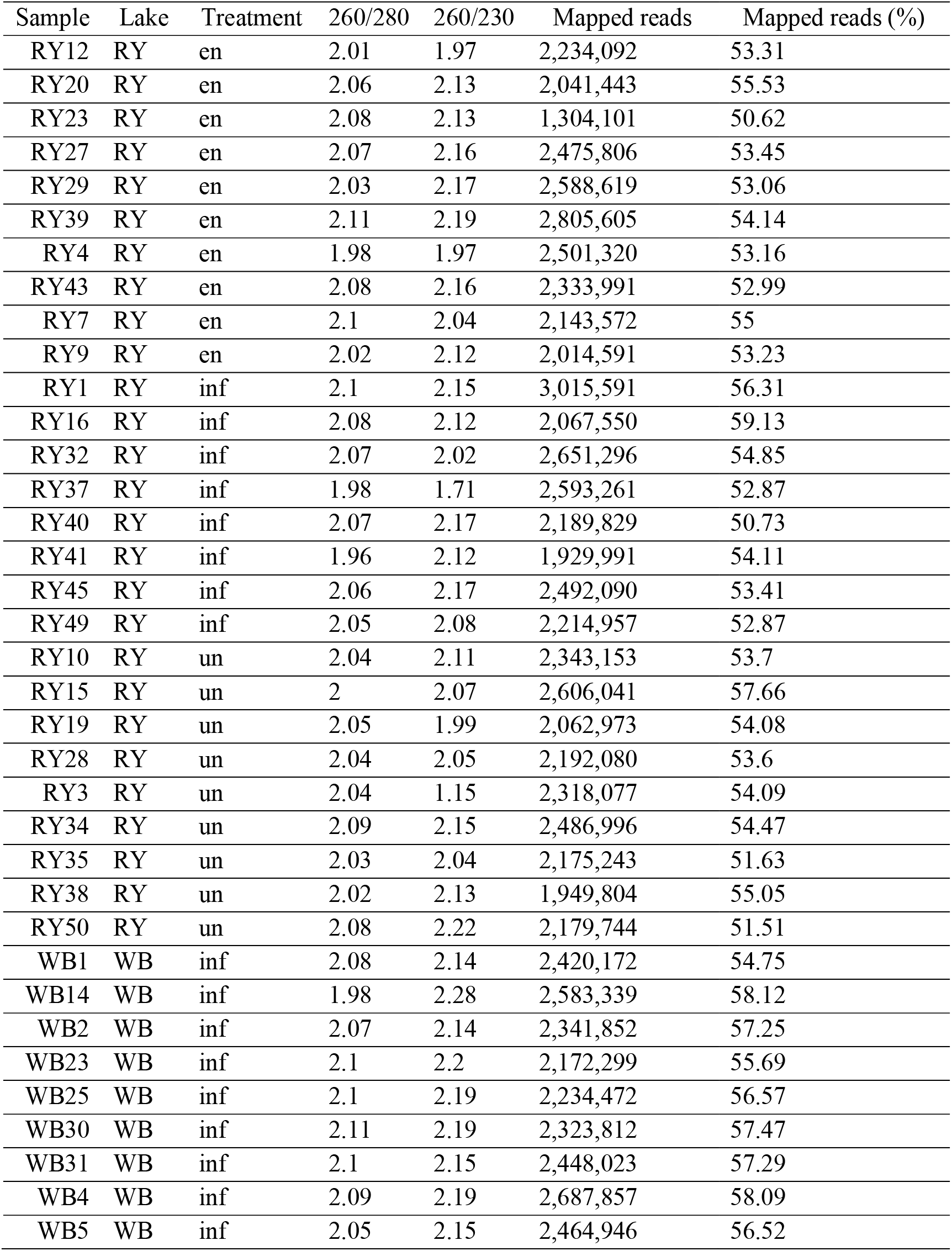
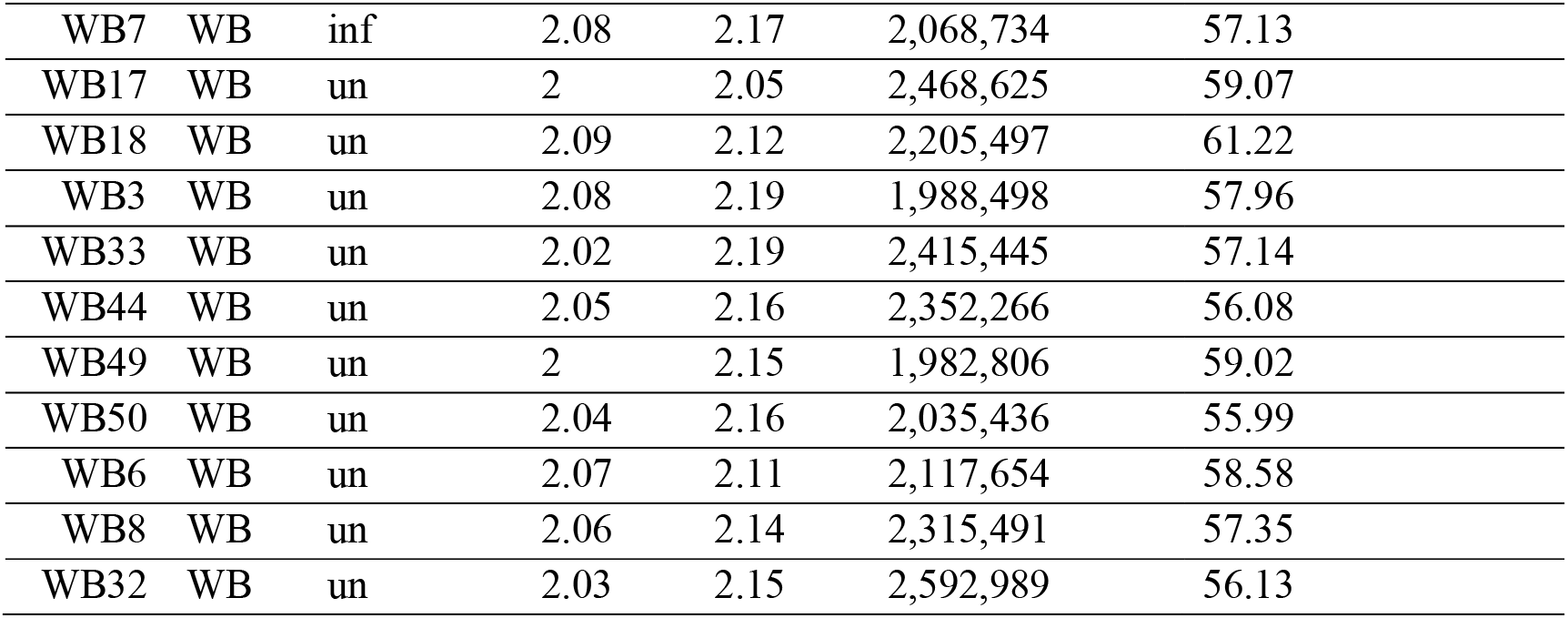
Quality Ratios (260/28 0 & 260/230) and alignment rates for each of the samples. en = stickleback with encysted S. solidus; inf = infected stickleback that did not encyst S. solidus; un = uninfected stickleback

**Table S2:**
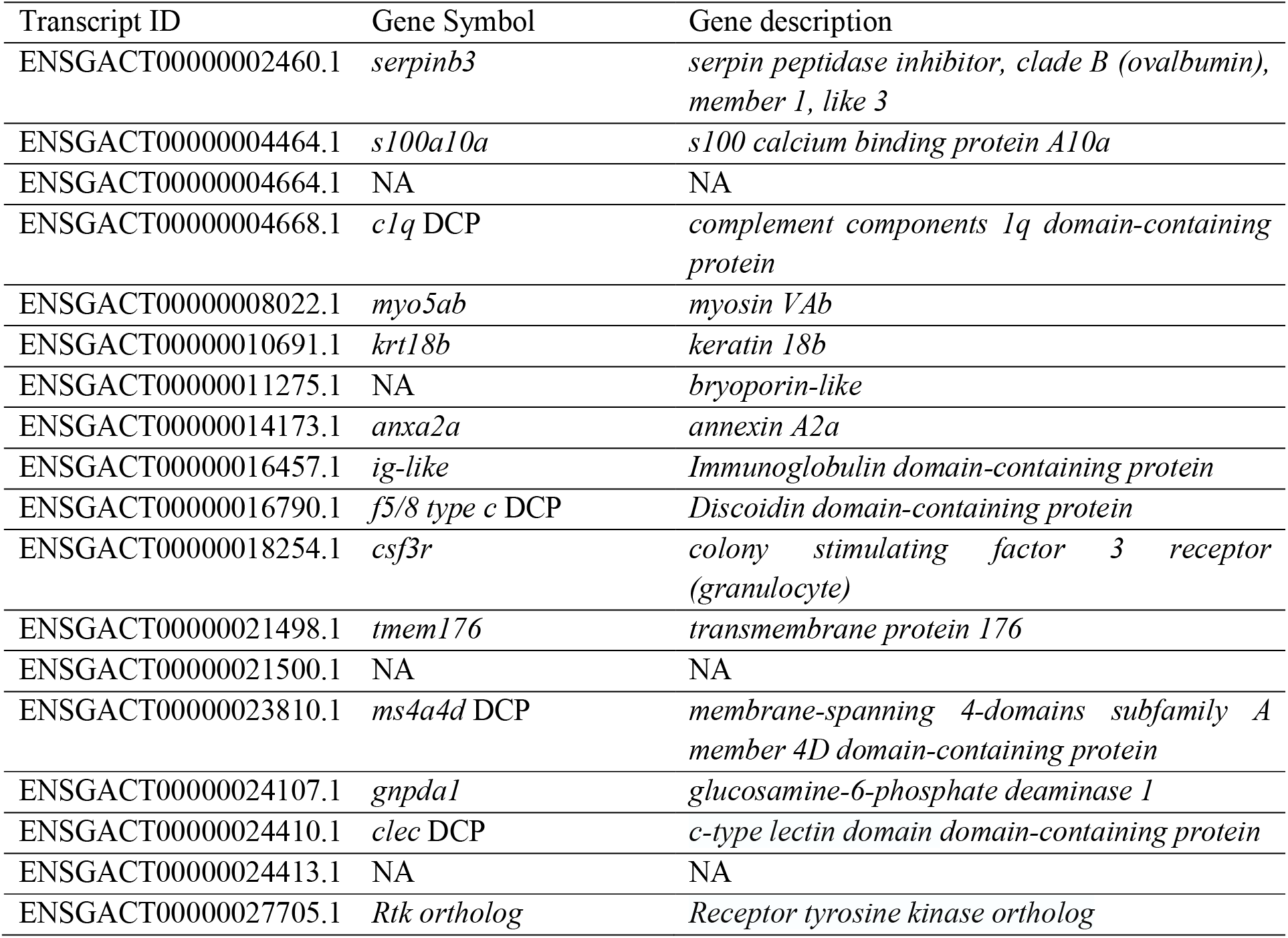
Transcript information associated with the black module for the Walby stickleback WGCNA. DCP = domain-containing protein

**Table S3:**
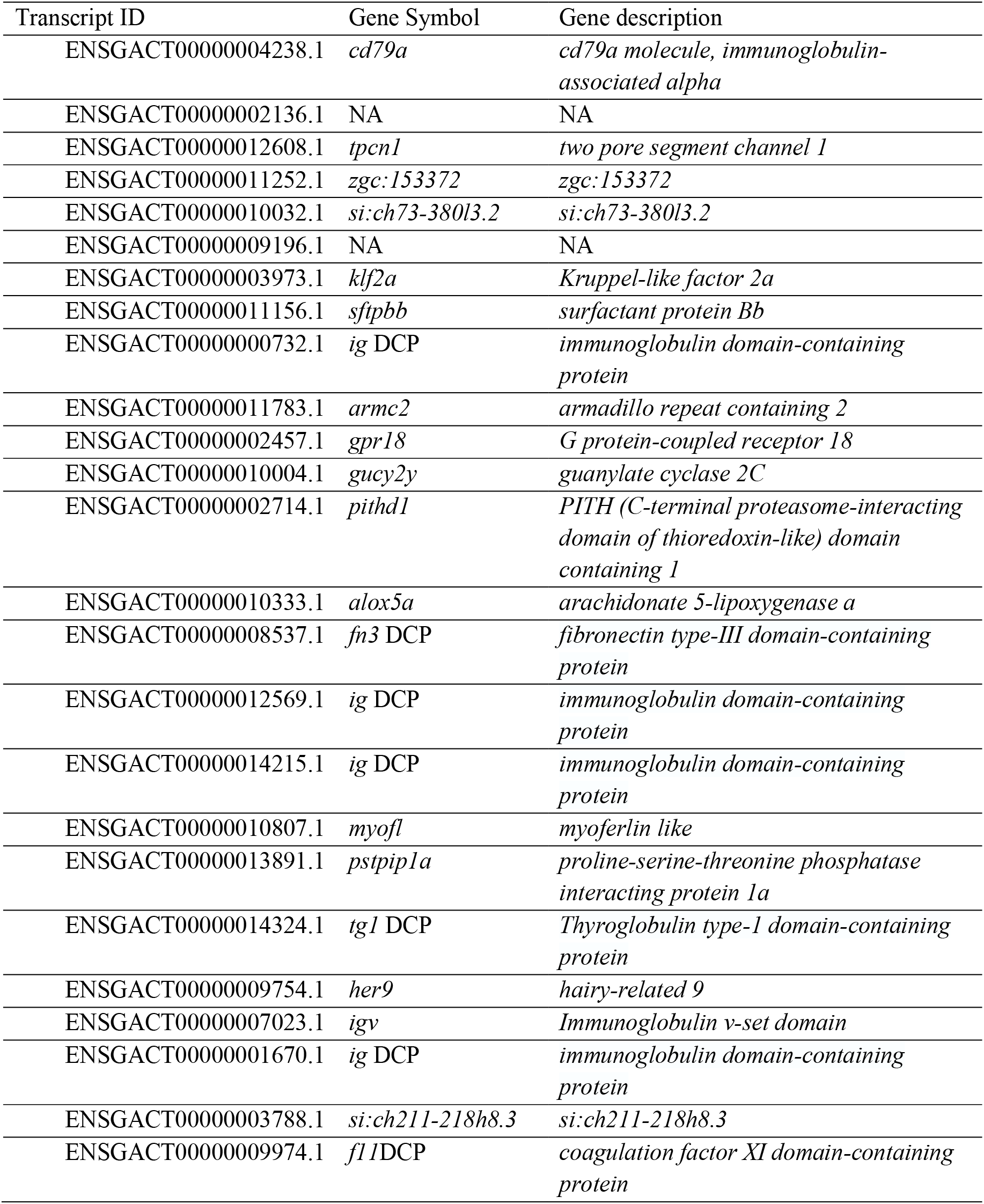
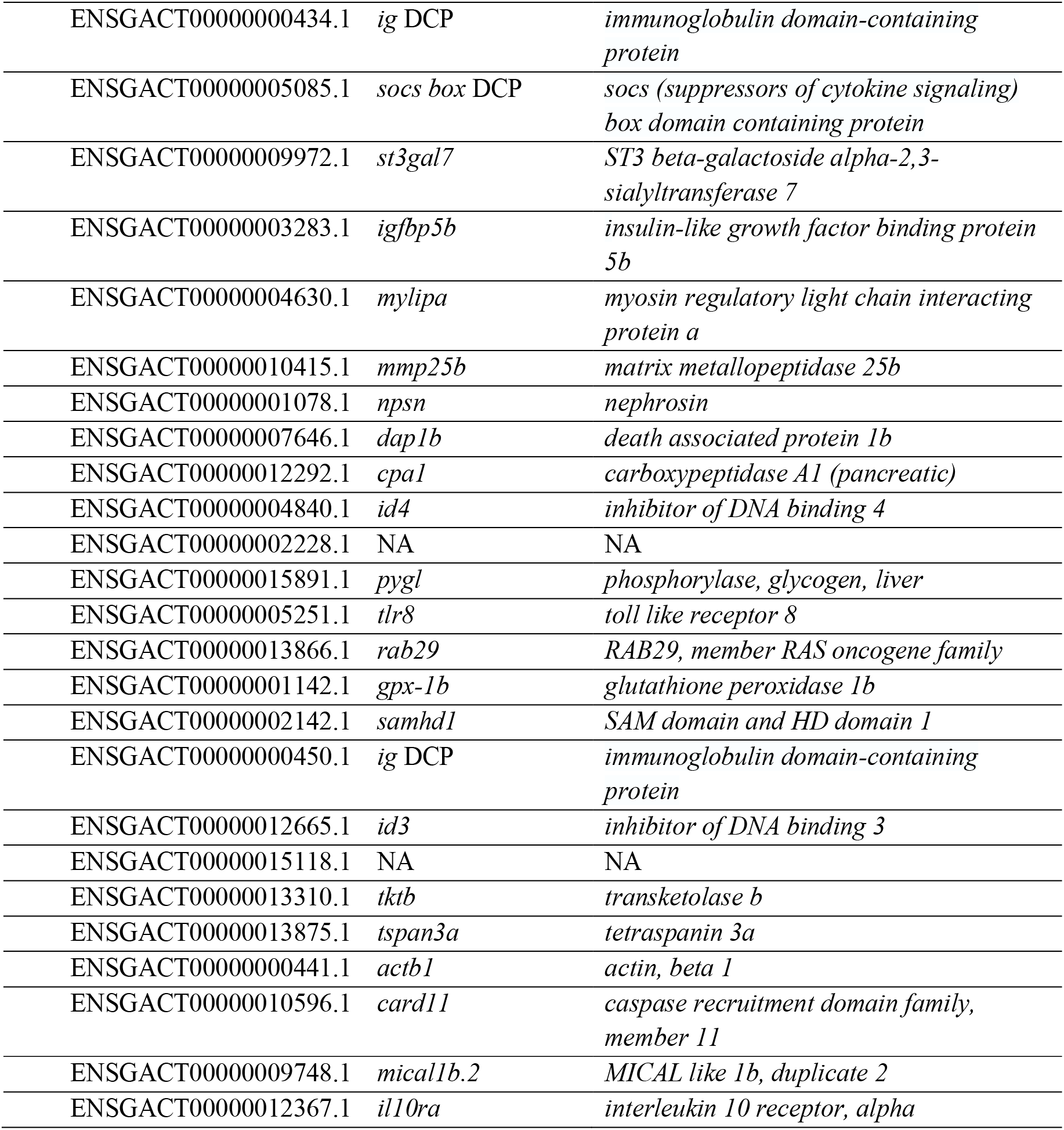
Transcript information associated with the brown module for the Walby stickleback WGCNA. DCP = domain-containing protein

**Table S4:**
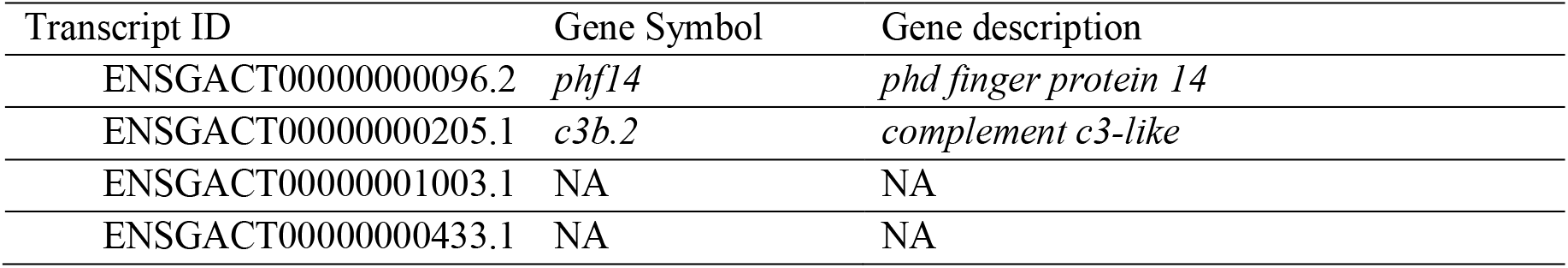
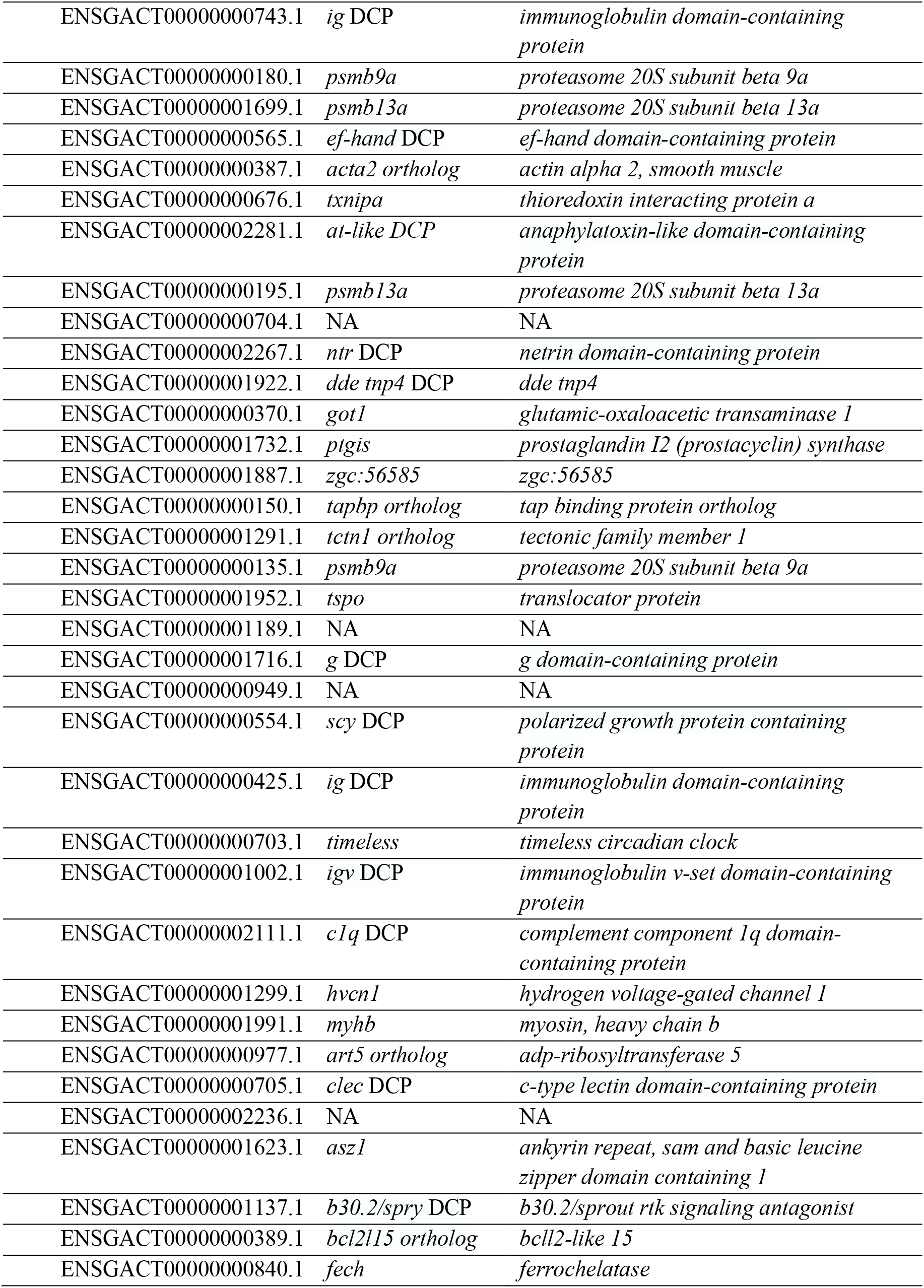
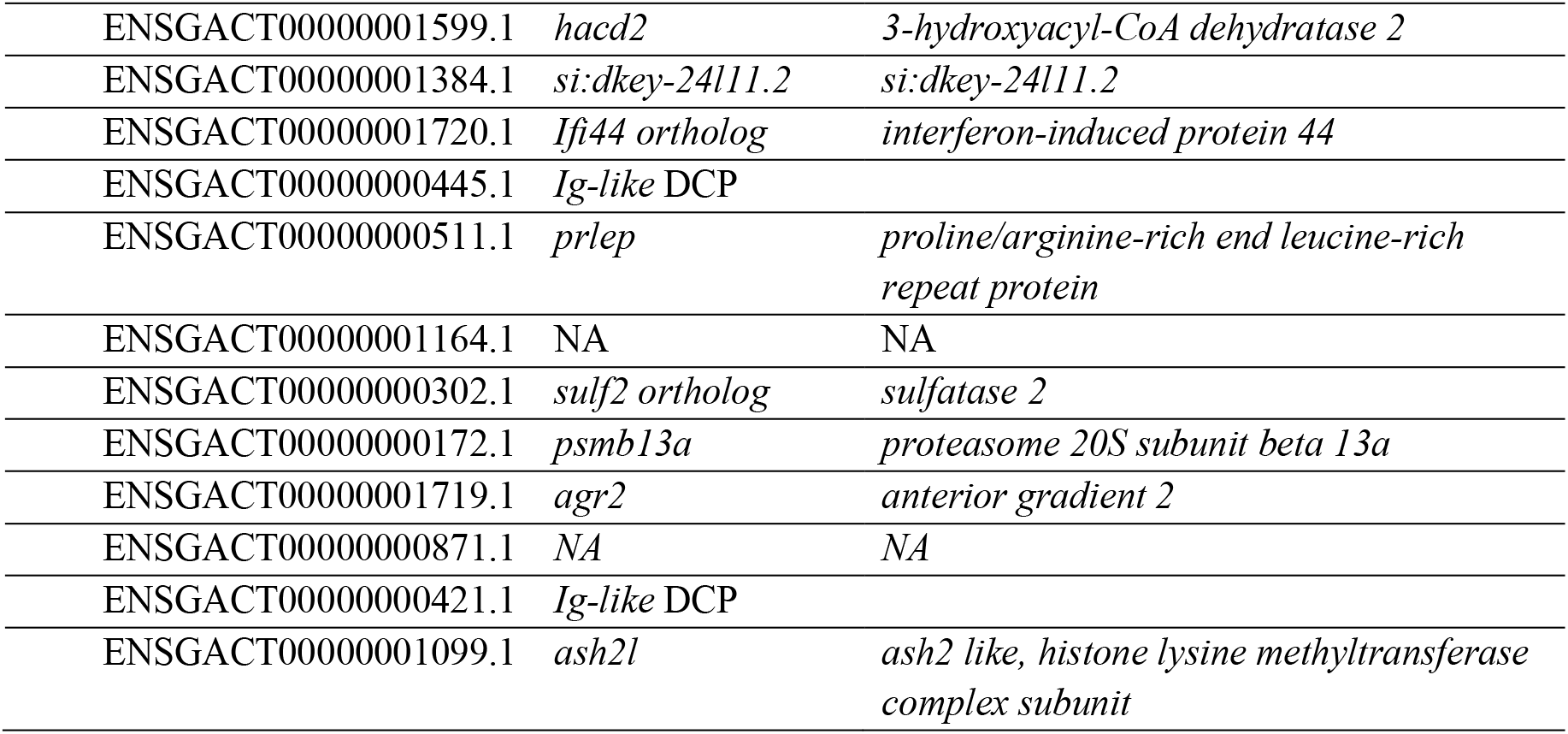
Transcript information associated with the grey module for the Walby stickleback WGCNA. DCP = domain-containing protein

**Table S5:**
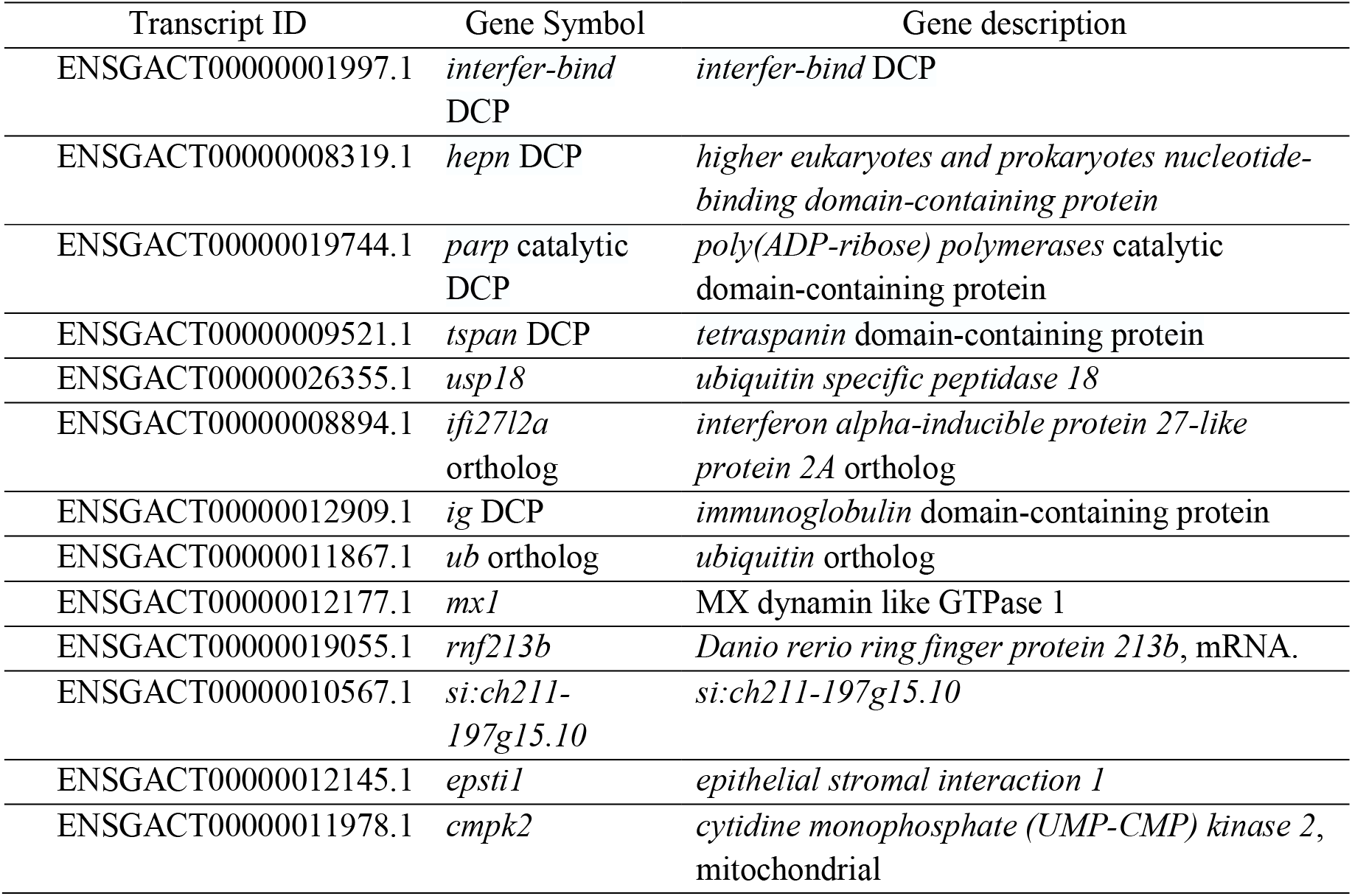
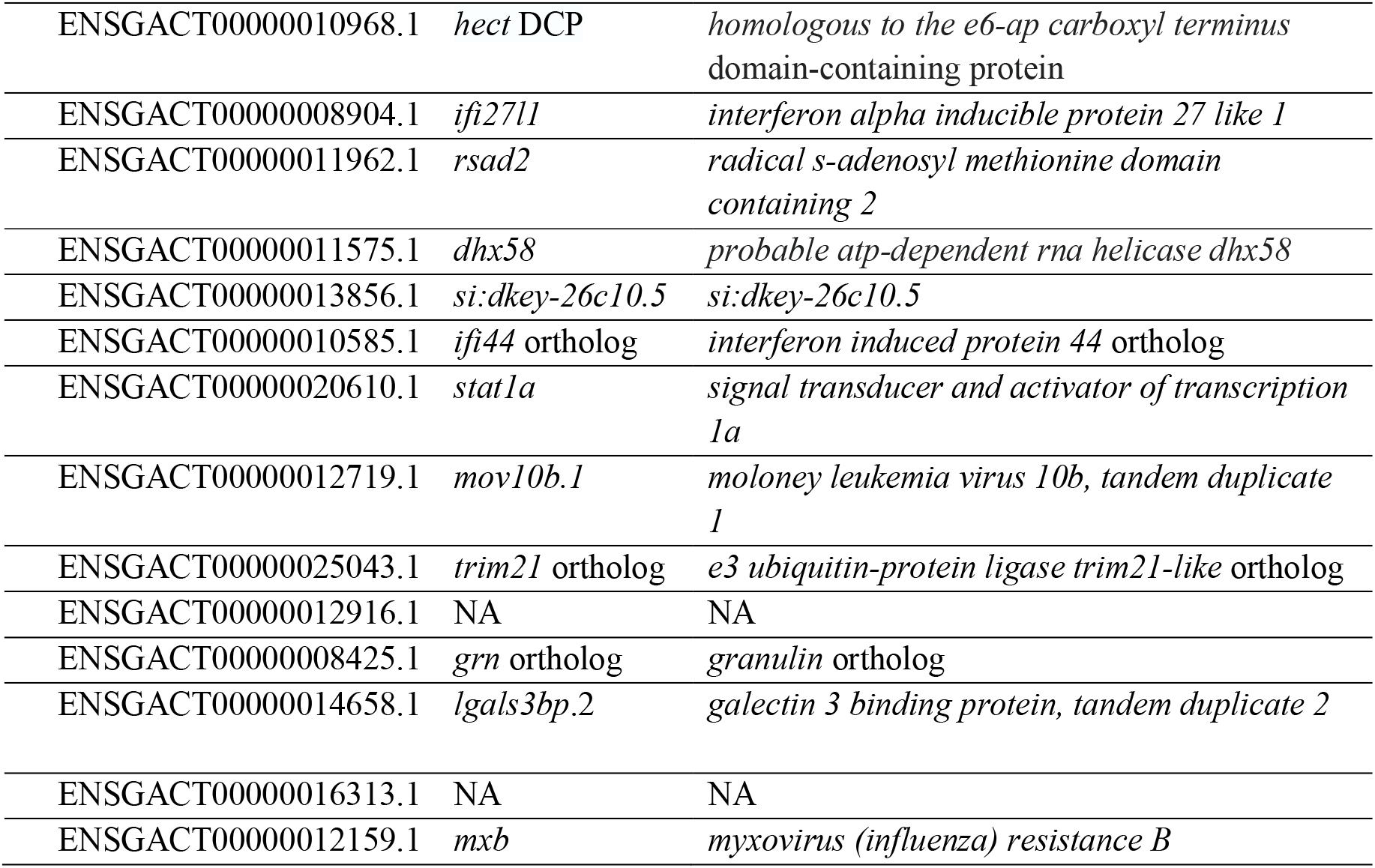
Transcript information associated with the red module for the Walby stickleback WGCNA. DCP = domain-containing protein

**Table S6:**
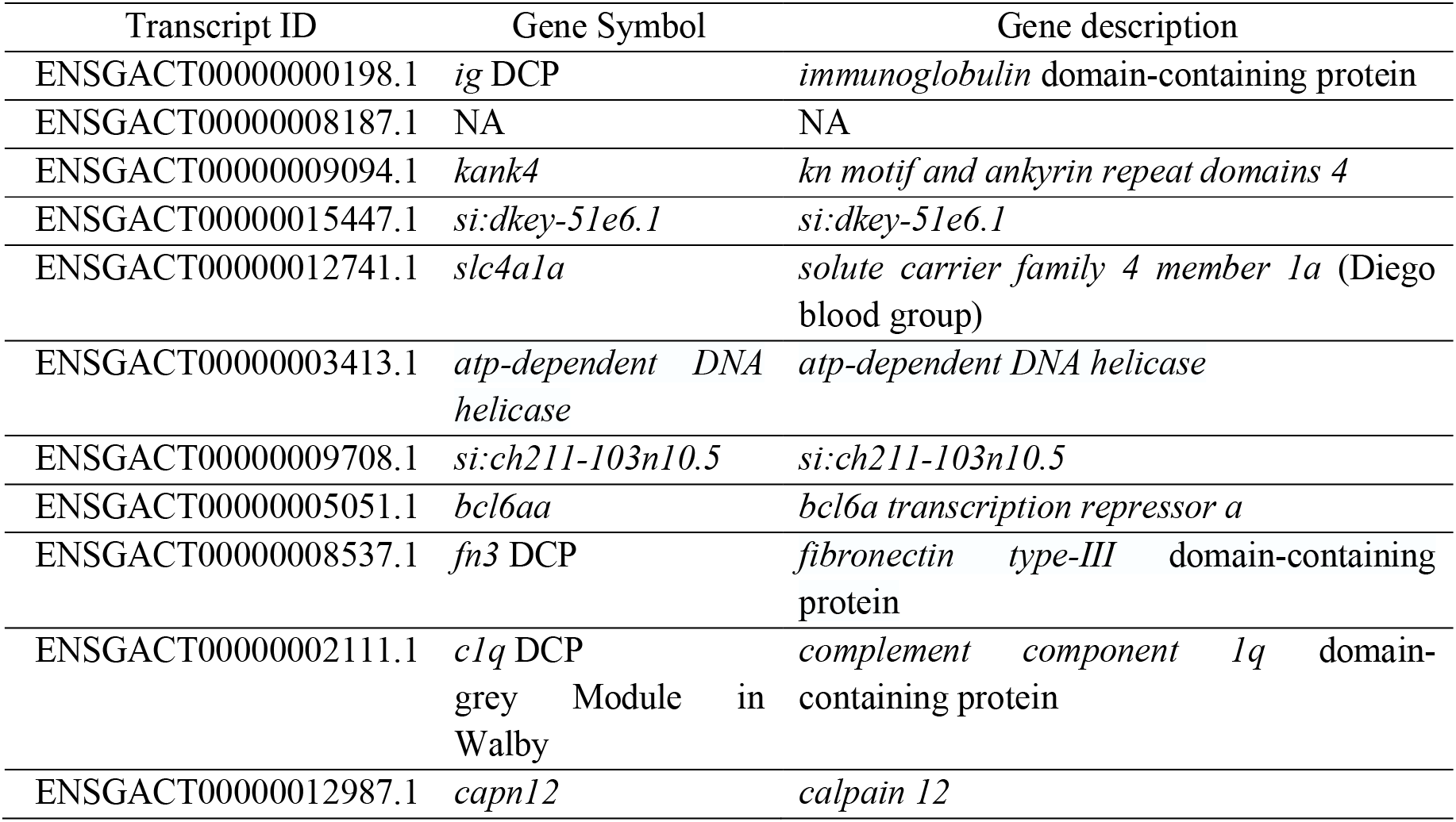
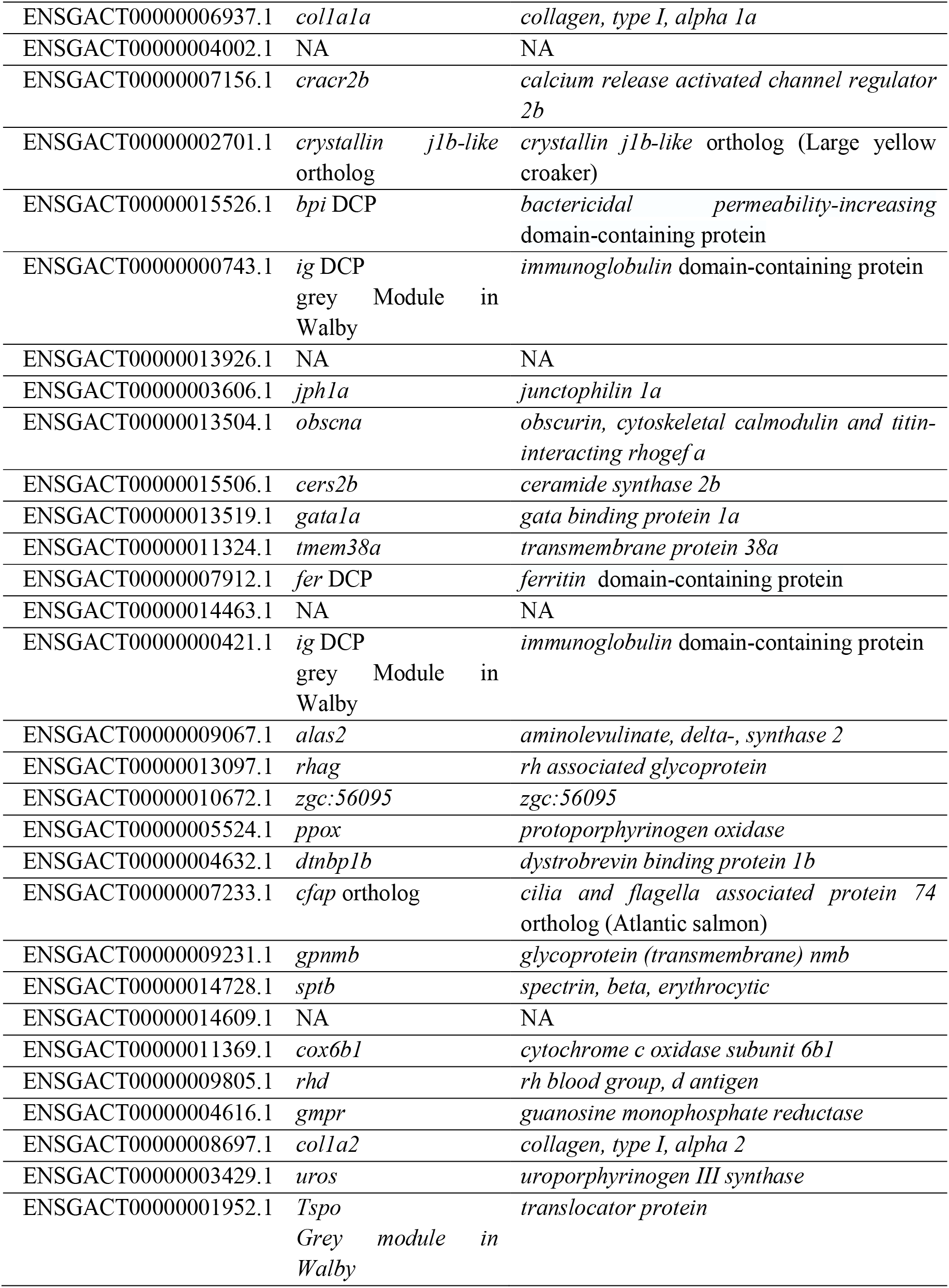
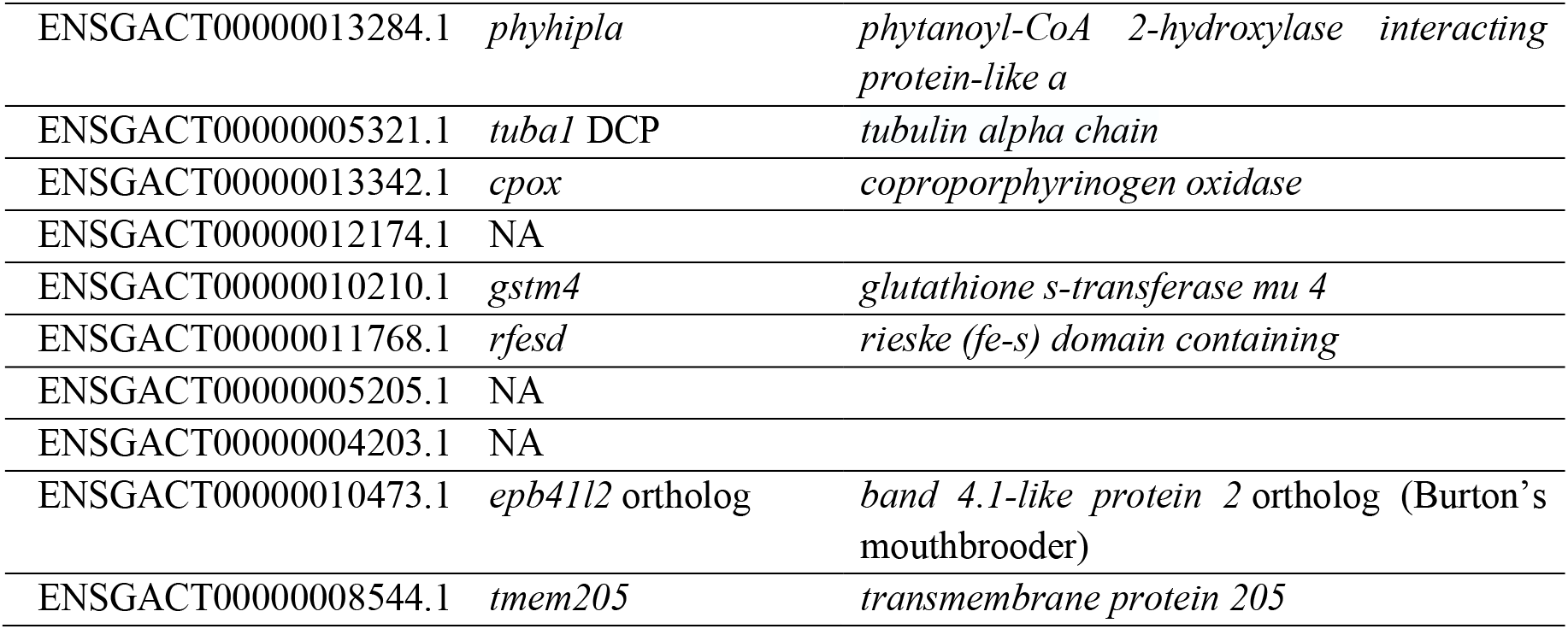
Transcript information associated with the green module for the Rocky Lake stickleback WGCNA. DCP = domain-containing protein

**Table S7:**
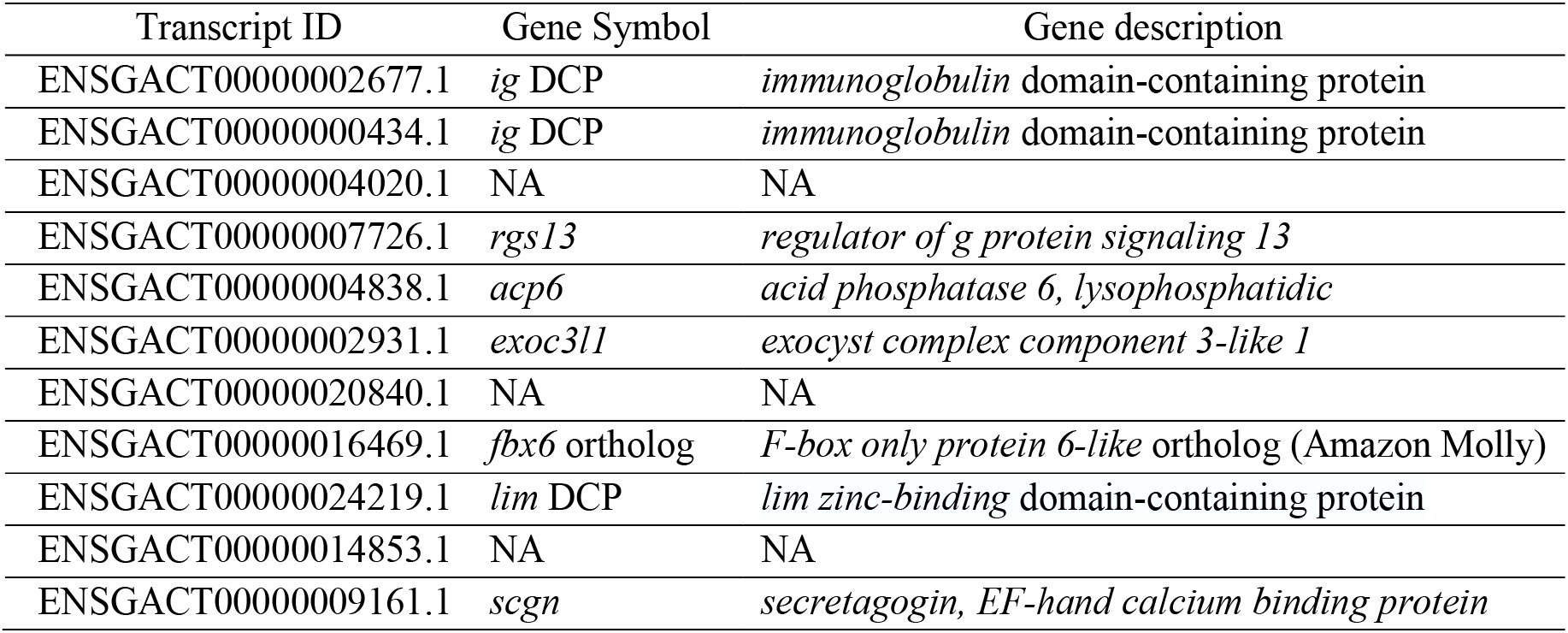
Transcript information associated with the grey module for the Rocky Lake stickleback WGCNA. DCP = domain-containing protein

| Transcript ID | Gene Symbol | Gene description |
| --- | --- | --- |
| ENSGACT00000002677.1 | *ig* DCP | *immunoglobulin domain-containing protein* |
| ENSGACT00000000434.1 | *ig* DCP | *immunoglobulin domain-containing protein* |
| ENSGACT00000004020.1 | NA | NA |
| ENSGACT00000007726.1 | *rgs13* | *regulator of g protein signaling 13* |
| ENSGACT00000004838.1 | *acp6* | *acid phosphatase 6, lysophosphatidic* |
| ENSGACT00000002931.1 | *exoc3l1* | *exocyst complex component 3-like 1* |
| ENSGACT00000020840.1 | NA | NA |
| ENSGACT00000016469.1 | *fbx6* ortholog | *F-box only protein 6-like* ortholog (Amazon Molly) |
| ENSGACT00000024219.1 | *lim* DCP | *lim zinc-binding domain-containing protein* |
| ENSGACT00000014853.1 | NA | NA |
| ENSGACT00000009161.1 | *scgn* | *secretagogin, EF-hand calcium binding protein* |

**Table S8:**
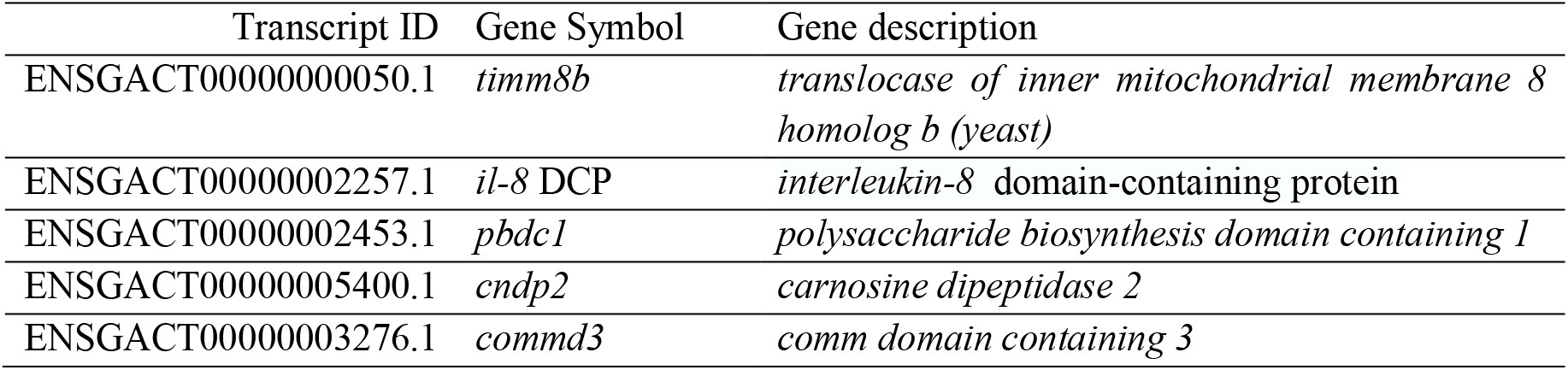
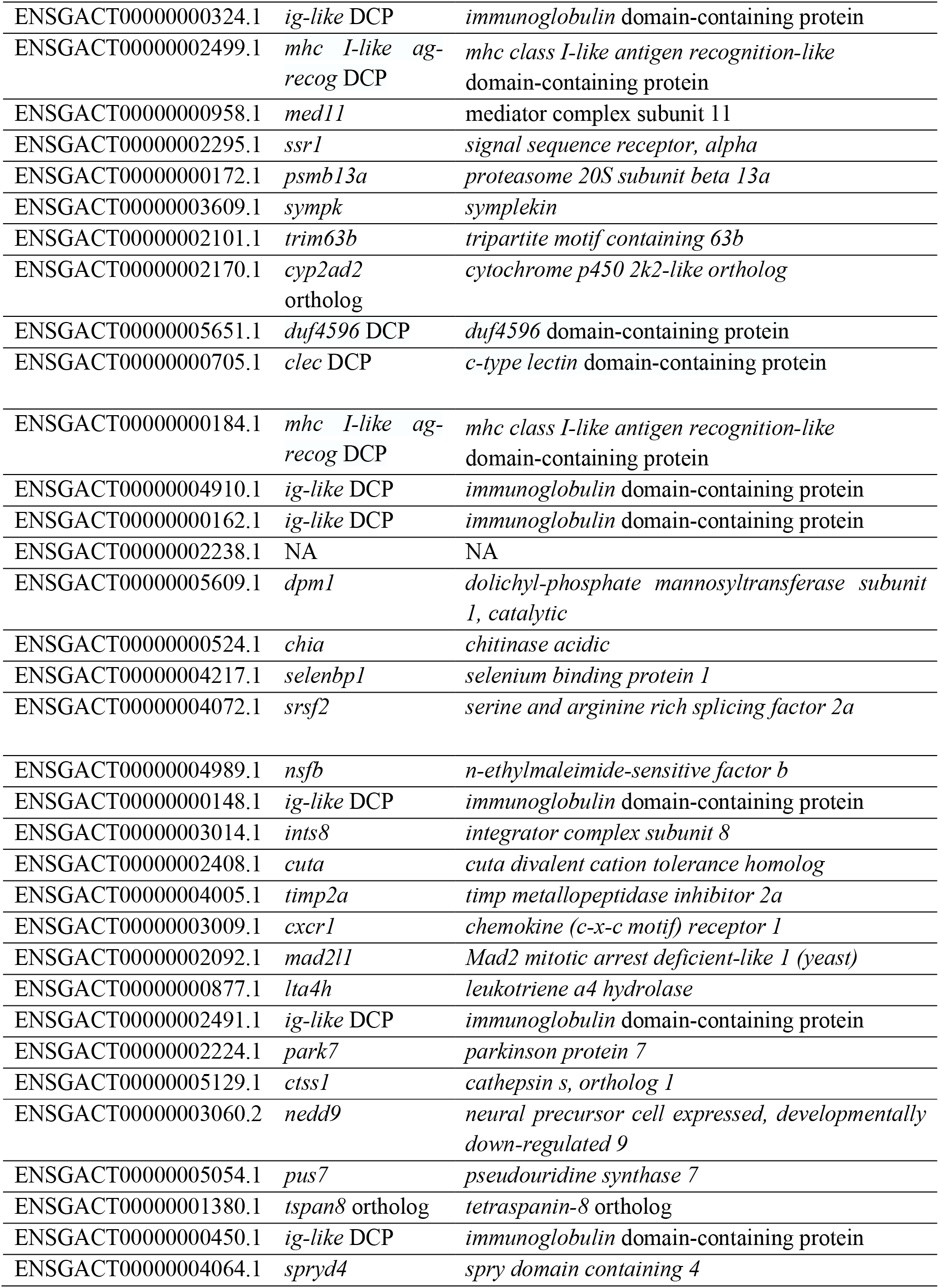
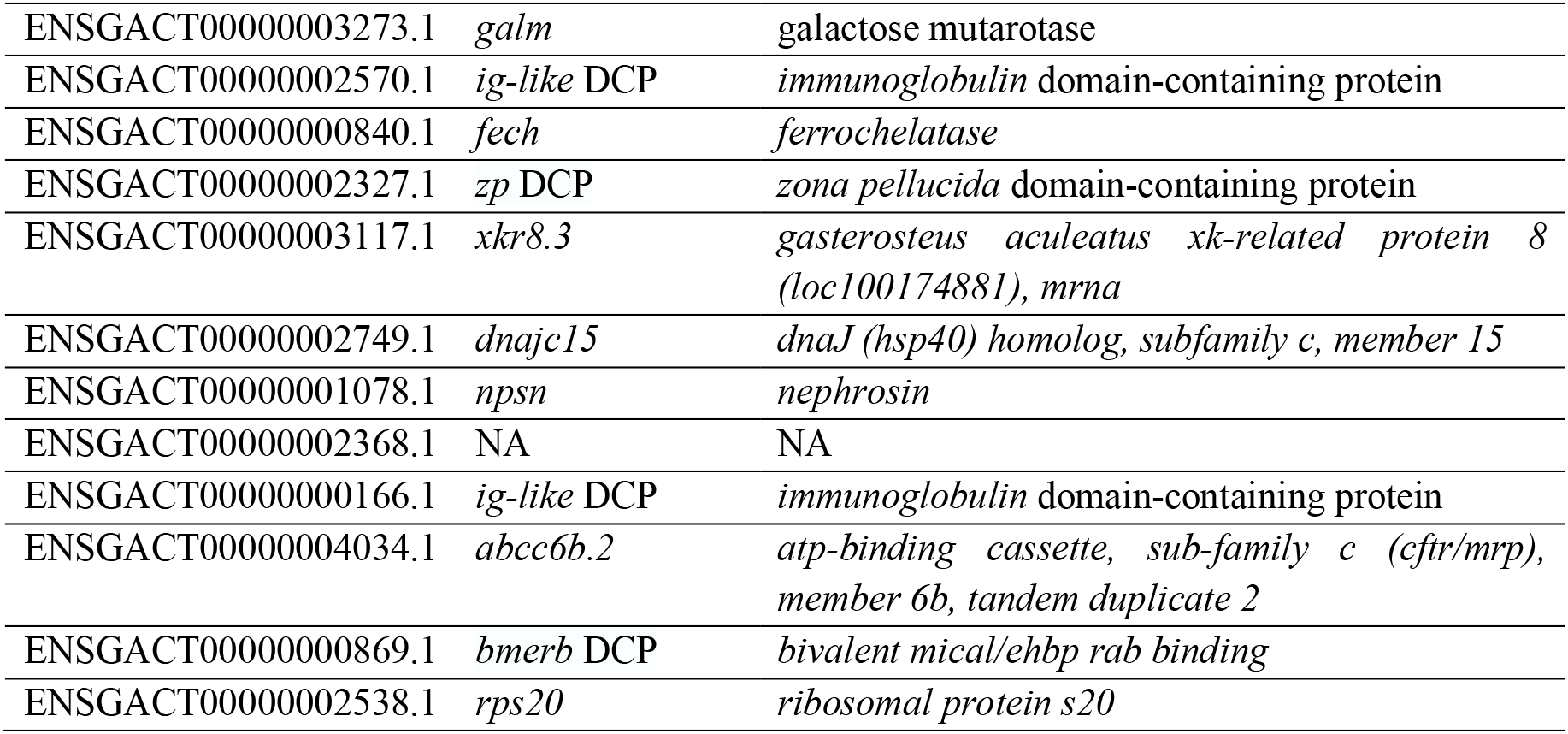
Transcript information associated with the blue module for the Rocky Lake stickleback WGCNA. DCP = domain-containing protein

